# Reversible haploidisation and convergent genomic routes to antifungal resistance in the *Candida parapsilosis* species complex

**DOI:** 10.64898/2026.05.19.725100

**Authors:** Juan Carlos Nunez-Rodriguez, Vladislav Biriukov, Alvaro Redondo-Rio, Mrinalini Parmar, Miquel Àngel Schikora-Tamarit, Islam Ahaik, Valentina del Olmo, Toni Gabaldón

## Abstract

Antifungal resistance is rising among non-albicans *Candida*, with outbreaks increasingly linked to drug-resistant *Candida parapsilosis*. Using large-scale experimental evolution under fluconazole and anidulafungin, coupled to phenotyping and genome sequencing, we define resistance strategies across *C. parapsilosis*, *Candida metapsilosis* and *Candida orthopsilosis*. Despite diverse genetic backgrounds, adaptation repeatedly converged on a shared genomic toolkit that combines protein-altering mutations in key regulators and drug targets (including MRR1 and FKS1) with copy-number changes, aneuploidy, as well as loss of heterozygosity driving resistance alleles to homozygosity. Strikingly, drug selection triggered recurrent but reversible haploidisation in *C. orthopsilosis*, revealing ploidy reduction as a transient, selectable survival strategy. Resistance, particularly after fluconazole selection, imposed fitness costs that promoted compensatory evolution and resistance loss after drug withdrawal. Together, antifungal adaptation in this complex affects convergent targets, while being plastic in its genomic routes, with implications for resistance surveillance and drug cycling therapies.

## Introduction

Invasive fungal infections cause more than two million deaths every year, and hospital-acquired invasive candidiasis remains a major clinical challenge^1,2^. Although *Candida albicans* still is the leading cause of candidiasis, infections caused by non-albicans *Candida* species are increasing, now collectively exceeding 50% prevalence^3^. Among these, *Candida parapsilosis* – typically the second to third most frequently reported *Candida* species– is of particular concern because of its intrinsic low echinocandin susceptibility, the growing incidence of fluconazole resistance, and the persistently high mortality despite active treatment^4^. These considerations, together with the increasing frequency of resistant outbreaks, led to its recent inclusion in the World Health Organization (WHO) list of high-priority fungal pathogens^5^.

*C. parapsilosis* belongs to the *Candida parapsilosis* species complex, which also includes the less prevalent pathogens *Candida orthopsilosis* and *Candida metapsilosis*^6^. While these species have indistinguishable morphologies, they are markedly different at the genomic level. *C. parapsilosis* is a largely homozygous diploid that exhibits low genetic variability compared to other *Candida* species^7–9^. In contrast, *C. metapsilosis* and *C. orthopsilosis* comprise phylogenetically complex populations dominated by highly heterozygous strains arising from multiple independent hybridisation events^10,11^. Members of this complex occur in diverse natural environments^6^, can colonize healthy individuals, and cause opportunistic infections that can be fatal in susceptible patients^12,13^.

Compared to *C. albicans, C. parapsilosis* exhibits a somewhat lower mortality (14.5% to 47%^14^), but is notable for its high transmissibility^15–18^ and its capacity to form robust biofilms on catheters and medical devices^19,20^, which facilitates persistence and spread in hospital environments. In contrast, the prevalence of *C. metapsilosis* and *C. orthopsilosis*, remains uncertain, partly due to frequent misidentification as *C. parapsilosis* in routine diagnostics^6,21^.

In recent years, infections caused by drug-resistant *C. parapsilosis* have increased, with multiple reports of hospital outbreaks associated with high mortality^22–27^. Given the limited drug armamentarium and the naturally low echinocandin susceptibility in this clade, the emergence of azole resistance is especially concerning. Resistance in the *C. parapsilosis* complex has been linked to mutations in canonical pathways—such as alterations in *FKS1* (and *ERG3*) for echinocandin resistance and changes affecting the ergosterol pathway and its regulators (*ERG11*, *MRR1*, *TAC1* or *UPC2*) for azole resistance^6,28,29^– yet, the broader repertoire and the genomic mechanisms resulting in resistance across the clade remain largely unexplored^30,31^. Beyond protein-altering point mutations and other small variants, copy number variations (CNVs), and whole or partial chromosome aneuploidies have also been reported as adaptive responses to drug exposure^32–34^. While aneuploidies and CNVs have received recent attention in *C. parapsilosis*^9,35–38^, their roles in *C. metapsilosis* and *C. orthopsilosis*, remain unexplored. Moreover, as hybrid lineages are highly heterozygous, loss of heterozygosity (LOH) may play a specific role in drug adaptation by rapidly fixing resistance alleles, as demonstrated for *C. albicans*^39,40^. However, this possibility has not been explored in the species complex.

Here we sought to obtain a systematic and genome-wide view of the mechanisms of antifungal adaptation across the *C. parapsilosis* complex using large-scale experimental evolution under fluconazole and anidulafungin selection, followed by genomic and phenotypic characterization of evolved lineages representing the phylogenetic breadth of the clade. This design enables high resolution identification of de novo resistance mechanisms by direct comparisons of evolved isolates with their closely related naive parentals, a comparison often unavailable when only considering clinical isolates^41,42^. Our experiment involved 288 evolving populations, over 10,0000 phenotypic measurements, and detailed genomic characterisation of 111 strains. Our results show that the species within the complex can rapidly develop resistance using conserved mechanisms, irrespective of their environmental or clinical origin. This adaptation is achieved through a shared genomic alteration toolkit combining point-mutations, aneuploidies, CNVs and LOH, and is generally accompanied by drug– and background-dependent fitness trade-offs that shape subsequent compensatory evolution.

Strikingly, we uncover haploidisation as an adaptive response to drug exposure in this clade. Together, our findings comprehensively define the genomic landscape of resistance evolution in this emerging pathogen complex and provide a framework for improving surveillance of antifungal resistance.

## Results and discussion

### Directed experimental evolution reveals high adaptive potential to antifungal drugs in the *C. parapsilosis* complex

To characterise antifungal adaptation and generate azole– and echinocandin-resistant strains representative of the diversity within the *C. parapsilosis* complex, we performed experimental evolution under antifungal selection with a panel of isolates of *C. parapsilosis*, *C. metapsilosis*, and *C. orthopsilosis* (**Figure 1A**, **Supplementary Table 1**). The panel covers the phylogenetic breadth of pathogenic species in this complex^6^, including clinical and environmental isolates, hybrids, and, when available, their homozygous parental lineages^11^.

**Figure 1.**
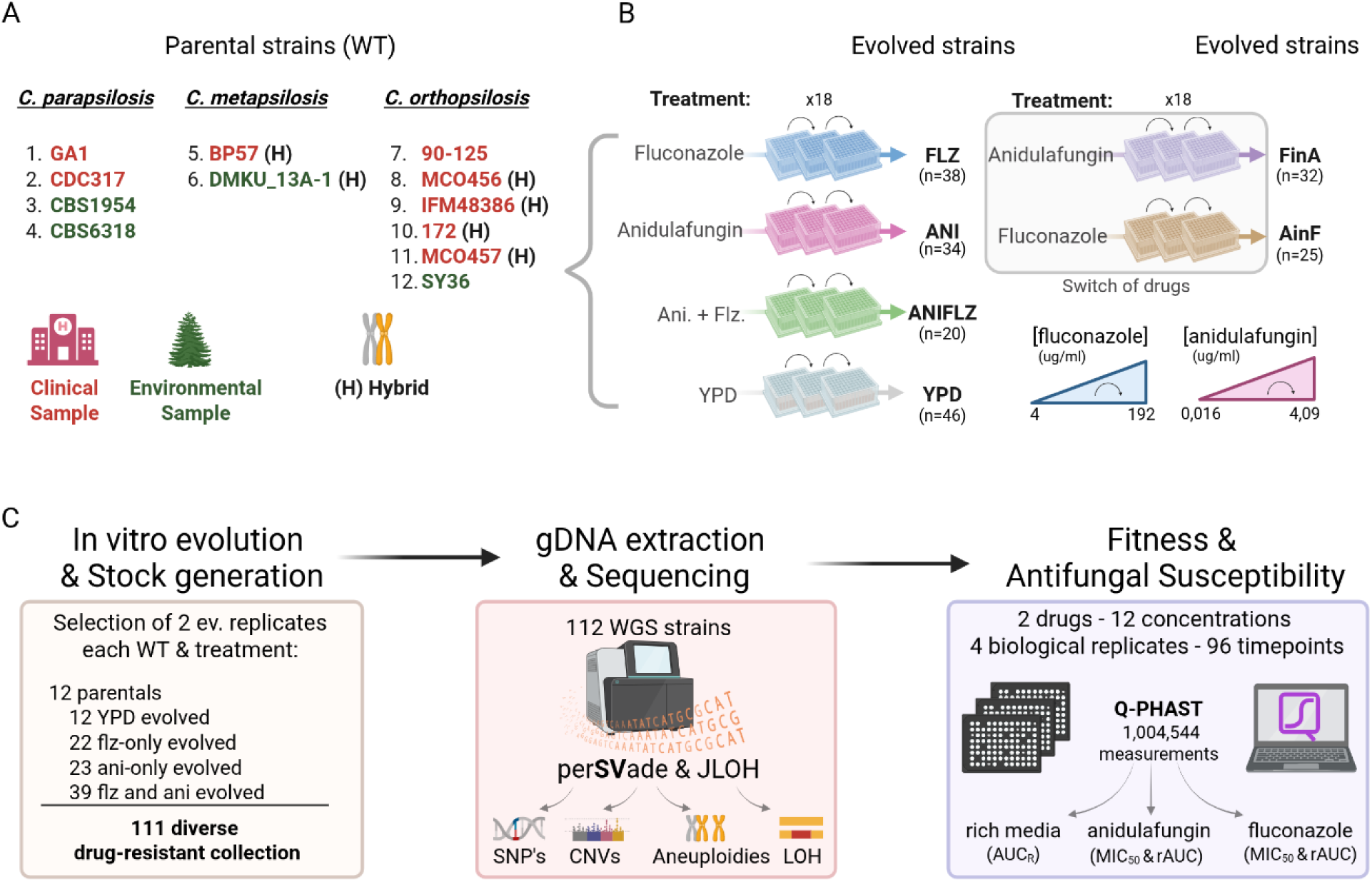
Experimental evolution and characterization. (**A**) List of wild type (WT) strains used. Red and green indicate the clinical or environmental origin of the strain, respectively. Hybrid strains are indicated with an (H) at the end of the name. (**B**) Experimental evolution design. Briefly, WTs were evolved for 18 passages in different drug regimes, with incremental concentrations of fluconazole (from 4 to 192 µg/ml) and/or anidulafungin (from 0.016 to 4.09 µg/ml), leading to the evolved strains FLZ, ANI, ANIFLZ and YPD. Then, a drug-switch treatment was performed, evolving FLZ strains in anidulafungin (FinA) and ANI strains in fluconazole (AinF). (**C**) After in vitro evolution and stock generation, 111 strains were selected for characterization of SNPs, CNV, aneuploidies and LOH by whole genome sequencing, and were phenotyped to assess fitness in rich medium (AUC_R_) and in 12 concentrations of anidulafungin and fluconazole (rAUC and MIC_50_).

We evolved WT strains under different drug regimes using a serial passaging approach adapted from Ksiezopolska et al. 2021^43^ (see Methods). In brief, we propagated four replicates per strain for 18 passages, while gradually increasing concentrations of fluconazole (FLZ treatment), anidulafungin (ANI treatment), or both drugs simultaneously (ANIFLZ treatment) (**Figure 1B**). To assess sequential therapy consequences, FLZ and ANI evolved strains were subsequently evolved under the alternative drug (FinA and AinF, respectively). In parallel, we evolved control populations in drug-free medium (YPD treatment). The experiment lasted approximately 120 days and comprised up to 288 evolving populations.

Most strains (195 strains, 73.58%) survived the whole experiment, at least in one of the replicates, indicating high adaptive capacity across the *C. parapsilosis* complex (**Supplementary Table 2**). Survival was high under monotherapy (38/48 and 34/48, for FLZ and ANI, respectively), but markedly reduced in the combined treatment (20/48, ANIFLZ), indicating that simultaneous targeting of distinct cellular processes constrained adaptation. In contrast, both drug-switch regimes exhibited similar survival percentages as monotherapy (25/35 and 32/38, for AinF and FinA, respectively), suggesting limited cross-resistance or collateral sensitivity between these two drugs^44^.

Compared with an equivalent study in *N.glabratus*^43^, where all populations survived monotherapy and 56% survived the combined treatment, our results suggest a lower adaptation ability to these two drugs in the *C. parapsilosis* complex. This is particularly true outside *C. parapsilosis*, with survival rates being 62.50% and 58.33% in *C. orthopsilosis* and *C. metapsilosis*, respectively, but 80.21% in *C. parapsilosis* (**Supplementary Table 2)**. Notably, no *C. metapsilosis* population survived ANIFLZ (**Supplementary Table 2**). Extinctions were concentrated in a subset of genetic backgrounds with 70% of extinct lineages deriving from five of twelve WTs (Co_90-125, Co_SY36, Co_MCO456, Cp_CBS6318 and Cm_BP57). Co_90-125 (parental A of *C. orthopsilosis* hybrids), was particularly vulnerable with only 33% surviving population, and none surviving FLZ and ANIFLZ treatments (**Supplementary Table 2**). Notably, *C. orthopsilosis* hybrids generally outperformed their parentals. These results support the hypothesis that hybridisation can increase the probability of adaptation in *C. orthopsilosis*, providing an advantage in the clinical setting as previously hypothesized^10,11^. Finally, we observed no significant differences in survival between environmental and clinical isolates.

Following experimental evolution, we selected two evolved isolates per WT and drug treatment for genomic and phenotypic characterisation. For each WT, we additionally included the naïve ancestor and one YPD-evolved replicate as controls. This totalled 111 strains undergoing phenotyping and whole-genome sequencing (**Figure 1C**). We quantified fitness and drug susceptibility using Q-PHAST^45^, resulting in >10,000 growth curves with 96 time points from which we computed AUC, rAUC, MIC_50_ and MIC_R parameters (see methods).

Adapted isolates from the three species reached comparable levels of drug resistance, with no significant differences between environmental and clinical strains, or between hybrids and their parental lineages, suggesting shared underlying mechanisms (All tests p-value > 0.05 after FDR correction, **Supplementary Figure 1A, B, Supplementary Tables 3-5, Supplementary Note 1**). All fluconazole-evolved strains (FLZ, ANIFLZ, and AinF) developed resistance to this drug. Notably, one anidulafungin-evolved lineage from the WT with the lowest survival (Co_ANI_7F from Co_90-125) displayed partial cross-resistance to fluconazole (MIC50 = 8 µg/ml), a cross-resistance pattern previously reported in *N glabratus*, where it affected ∼50% of anidulafungin-evolved strains^43^.

In anidulafungin-exposed treatments (ANI, ANIFLZ, FinA), most isolates developed anidulafungin resistance (35/49, Supplementary Figure 1B; Supplementary Tables 3–5). However, in contrast to fluconazole, a subset of survivors either remained susceptible or grew insufficiently for robust susceptibility estimation, particularly in the combined ANIFLZ condition, in which no strain developed multidrug resistance (Supplementary Figure 1B). This finding was surprising given that these strains had successfully survived higher anidulafungin concentrations during the experimental evolution.

To investigate how strains classified as anidulafungin-non-resistant nevertheless survived selection, we re-tested fourteen anidulafungin-susceptible survivors using broth microdilution (at 24 and 48 hours), spot assays, and recovery of viable colonies from 4 µg/ml anidulafungin (**Supplementary Figure 2A–D**). Only one survivor showed a clear MIC increase consistent with acquisition of resistance, whereas four exhibited increased delayed growth or supra-MIC growth consistent with tolerance. Strikingly, the remaining isolates produced abundant viable colonies after prolonged exposure to 4 µg/ml, despite lacking a detectable and stable resistance phenotype by standard MIC criteria. Moreover, several WT backgrounds also retained viability at supra-MIC concentrations, indicating an intrinsic strain– and species-specific ability to withstand supra-MIC echinocandin exposure.

Overall, these results clearly show that antifungal exposure drives resistance acquisition in all strains and species of the *C. parapsilosis* complex. Combined ANIFLZ treatment increases extinction and prevents the simultaneous emergence of multidrug resistance, while sequential treatment has the same extinction ratio as monotherapy and allows strains to develop multiresistance. Remarkably, we demonstrated that evolved non-resistant strains survived treatment with supra-MIC concentrations of anidulafungin. Importantly, the observation of supra-MIC survival in isolates lacking stable resistance suggests that natural and/or acquired tolerance mechanisms may contribute to therapeutic failure by enabling persistence and regrowth after treatment discontinuation, even when routine testing does not detect resistance.

### Hotspot mutations in *MRR1* dominate fluconazole resistance

To elucidate genetic mechanisms driving acquired resistance, we whole-genome sequenced the 111 phenotyped strains (**Figure 1C**). We identified variants acquired during adaptation by comparing genomes of drug-evolved strains with their respective WT and YPD-evolved counterparts. We prioritized genes that independently acquired mutations in different genetic backgrounds, as recurrent targets are more likely to underlie adaptation (**Figure 2A, B)**, but we report all identified variants **(Supplementary Tables 3-5**).

**Figure 2.**
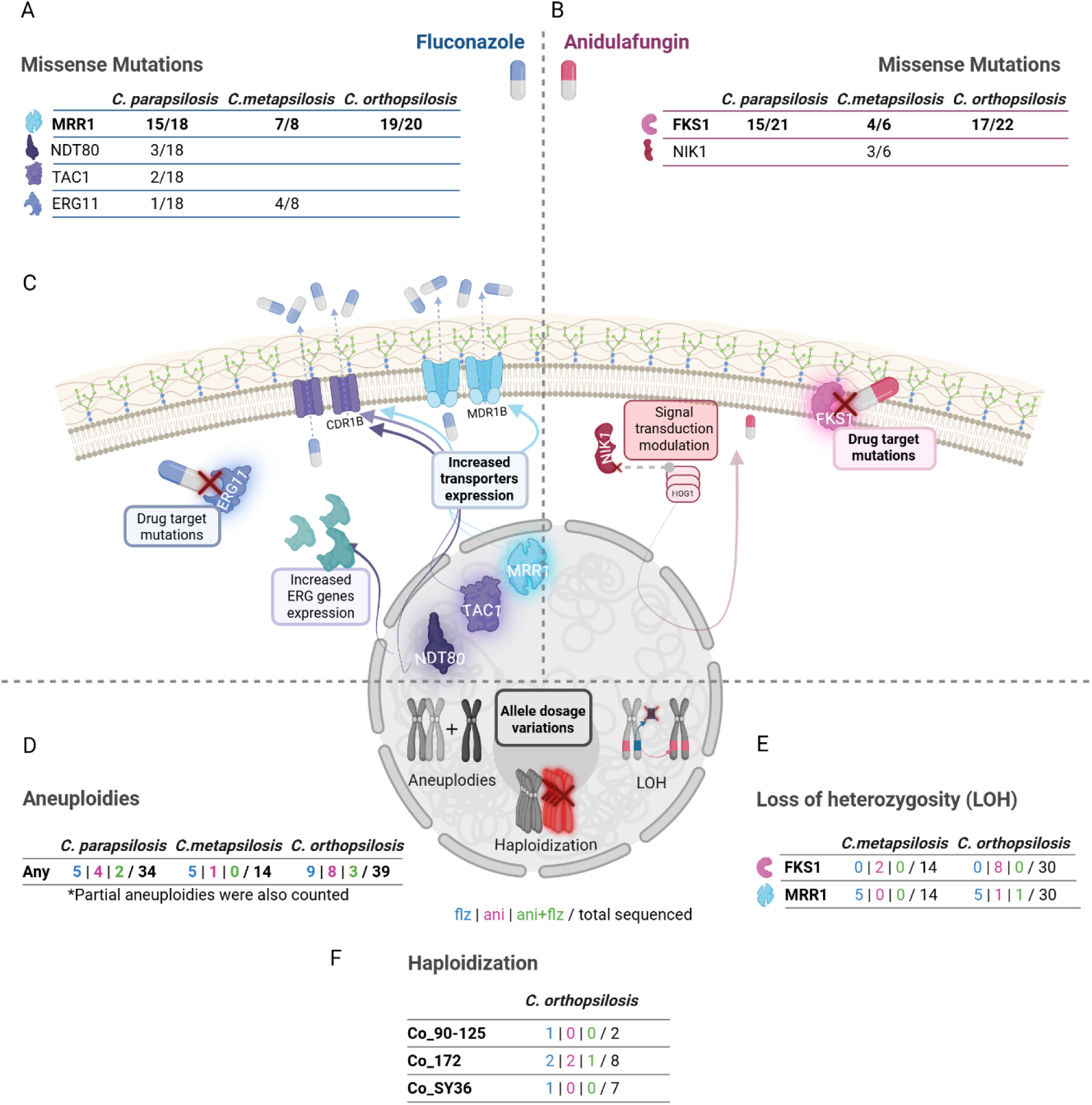
Main genomic modifications in resistant strains and its t of action. (**A-B**) Missense mutations in recurrently affected genes among *C. parapsilosis* complex strains after adaptation to **(A)** fluconazole or **(B)** anidulafungin. **(C)** Molecular pathways potentially linked to resistance, highlighting key proteins and processes identified in this study. **(D-F)** Count of **(D**) aneuploidies, **(E**) LOH or **(F)** haploidisation events observed under fluconazole (in blue) | anidulafungin (in pink) and anidulafungin plus fluconazole (in green) treatments versus the total of events counted in the corresponding strains sequenced.

Across fluconazole-evolved strains (FLZ, ANIFLZ, AinF), nearly 90% carried one of 45 missense mutations in *MRR1*. We established positional homology across the three species (**Figure 3A)** and compared these variants to 21 previously reported mutations (**Supplementary Table 6)**. Nine mutations detected in this study affect previously described positions^38,46^, with three matching the exact substitution: G604R^47,48^ in *C. parapsilosis* and two homologous variants in *C. orthopsilosis* (G292E and G581R)^38,47^. These overlaps support the clinical relevance of mutations identified through in vitro evolution.

**Figure 3.**
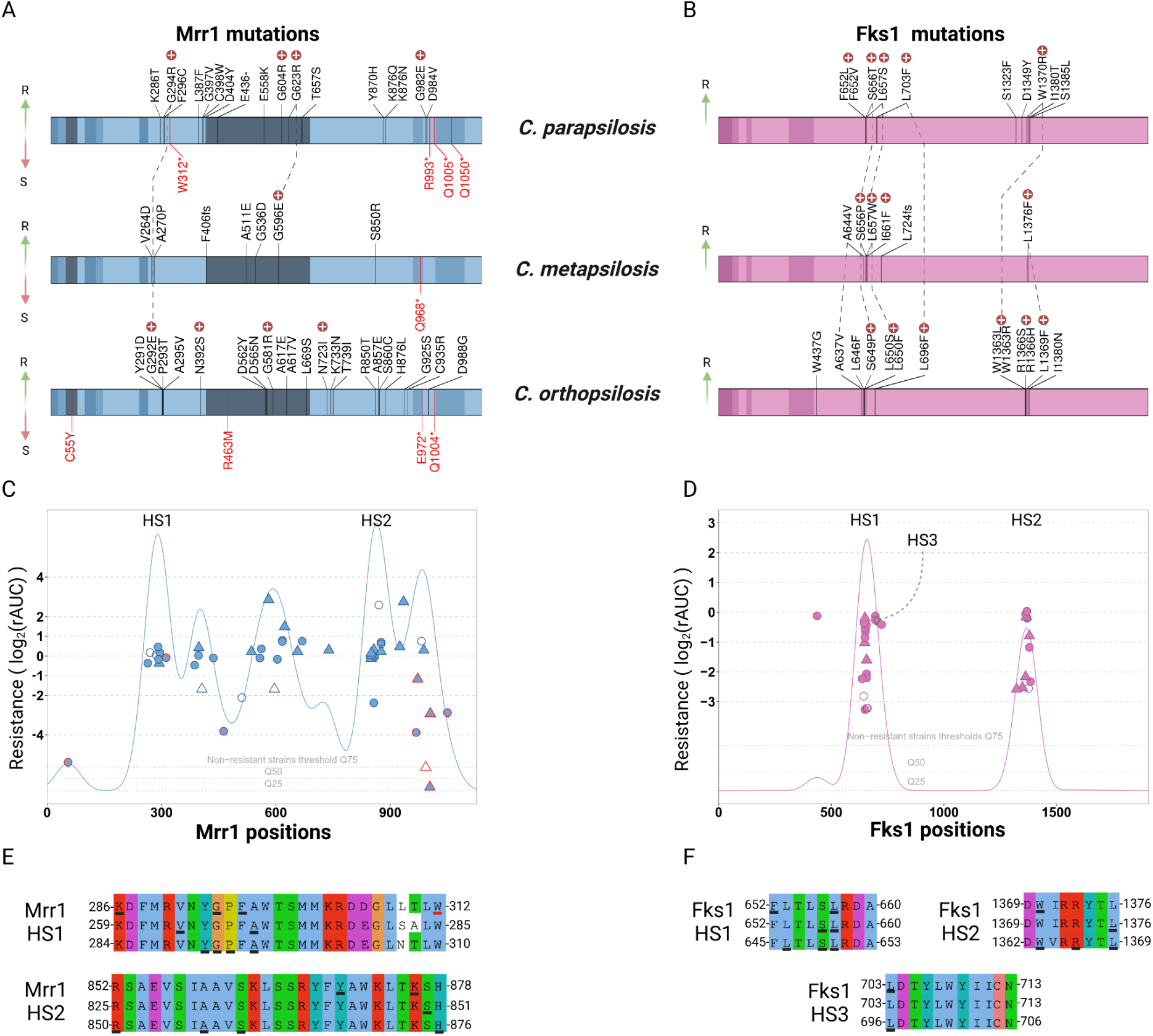
Mutations and hotspots in Mrr1 and Fks1. Identified mutations in Mrr1 **(A)** and Fks1 **(B)**. Shaded blocks show Pfam-predicted functional domains. Mutations at homologous positions between the three species are connected with dotted lines. Mutations above the bar are linked to an increase in resistance (R), while mutations below the bar (in red) are linked to susceptibility (S). Mutations marked with an encircled cross indicate positions of variants previously reported in clinical isolates (see **Supplementary Table 6** for details). (**C, D**) Position of mutations across Mrr1 and Fks1 proteins relative to the susceptibility of the strain, measured as log2 of the rAUC. Circles represent strains without aneuploidy, while triangles indicate strains with aneuploidy. Filled shapes denote strains with additional mutations in relevant genes (indicated in Figure 3 **A, B**). Shapes with a red border indicate compensatory mutations identified in this study. (**E, F**) Proposed hotspot regions in Mrr1 and Fks1, indicating homologous sequences in *C. parapsilosis*, *C. metapsilosis*, and *C. orthopsilosis*, respectively. Mutated residues in our experiments appear underlined in black, while positions with stop or compensatory mutation falling within hotspots appear in red.

In *C. albicans*, *MRR1* mutational hotspots have been defined in three protein regions^49^, and in *C. parapsilosis*, gain-of-function (GoF) variants associated with *MDR1B/CDR1B* overexpression cluster in the region encoding amino acids 852-875, which corresponds to the 873-896 hotspot in *C. albicans*^47,49,50^. In our experiment, *MRR1* mutations were concentrated in two regions: 852-878, confirming the previously described GoF hotspot, and 286-312, suggesting an additional hotspot in *C. parapsilosis* (**Figure 3C,E**).

Notably, during FinA switch conditions, we identified nine *MRR1* mutations that are likely compensatory as they appeared after loss of fluconazole selection and were associated with a decrease in fluconazole resistance (**Figure 3A,C,E, Figure 6D**, **Supplementary Tables 6**). These included premature stop codons and missense mutations predominantly (6/9 mutations) clustering within the region corresponding to the inhibitory domain in *C. albicans*^51^, but also affecting Zn(II)-C6 fungal type DNA binding (positions 38-71 in *C. parapsilosis*) and fungal specific transcription factor (420-693) domains. To our knowledge, this represents the first evidence for compensatory evolution in *MRR1* within the *C. parapsilosis* complex.

Beyond *MRR1*, we identified mutations in other transcription factors previously linked to azole resistance (*TAC1*, *NTD80*, and *UPC2*) but only in *C. parapsilosis* (**Figure 2A and Supplementary Table 3**). *TAC1* regulates *CDR* efflux pumps, and is a widely established fluconazole resistance mechanism in *C. parapsilosis*^23,29,52,53^. *NTD80* simultaneously regulates *CDR* pumps and ergosterol biosynthesis, and confers azole resistance in *C. albicans*^54^. In *C. parapsilosis*, *NTD80* and *UPC2* regulate some *ERG* genes but not *ERG11*^55,52,55^.

Mutations in *ERG11* showed a marked species bias: they were common in *C. metapsilosis* (4/8 fluconazole-evolved strains, always alongside *MRR1* mutations), rare in *C. parapsilosis* (1/18), and absent from *C. orthopsilosis* (**Supplementary Tables 3-5)**. *ERG11* encodes lanosterol 14-α-demethylase, an essential enzyme in ergosterol biosynthesis and the main target of azoles. Inhibition of *ERG11* leads to accumulation of toxic sterol intermediates^27,53,56^. While the evolved strain Cm_FLZ_6E harbored the Y132F substitution, a well-documented clinical variant associated with azole resistance^29,38,57–59^, the other four mutants (Cp_FLZ_4C, Cm_FLZ_5F, Cm_FLZ_6A, and Cm_AinF_6G) shared a novel I465F substitution. The recurrent appearance of I465F substitution strongly supports a role in azole resistance. Consistently, a recent study demonstrated that the equivalent substitution in *C. albicans* (I471F) is beneficial in the presence of fluconazole, clotrimazole, and voriconazole, but deleterious in isavuconazole, posaconazole or in the absence of antifungal, highlighting a context-dependent impact^60^.

Only 5 of 46 fluconazole-evolved strains (Cp_FLZ_2G, Cp_FLZ_4C, Cp_FLZ_4G, Cm_FLZ_6E and Co_AinF_7F) lacked *MRR1* mutations, implying alternative routes to resistance (**Supplementary Tables 3-5**). The three *C. parapsilosis* strains without *MRR1* mutations exhibited lower levels of fluconazole resistance as compared to other fluconazole-evolved strains, indicating that *MRR1* mutations confer stronger resistance and explaining their higher prevalence. Cp_FLZ_2G and Cp_FLZ_4G carried *TAC1* mutations, while Cp_FLZ_4C carried an *ERG11* mutation. Additionally, Cp_FLZ_4C and Cp_FLZ_4G, both derived from the same Cp_CBS6318 background, had mutations in *TAF12*, whose orthologs have been implicated in oxidative stress resistance in *C. albicans* (*TAF12L*)^61^, and in antifungal resistance in *Saccharomyces* quantitative trait loci (QTLs) studies^62^. In *C. metapsilosis*, the mutant carrying *ERG11* Y132F but lacking MRR1 mutation (Cm_FLZ_6E) exhibited the lowest resistance among evolved strains, despite the known clinical relevance of Y132F, underscoring that phenotypic outcomes can be strongly shaped by genetic background.

### A broad spectrum of *FKS1* variants mediates anidulafungin resistance

Analysis of anidulafungin-evolved strains identified mutations in *FKS1* as the predominant mechanism of echinocandin resistance across all three species (**Figure 2B**). Of the 50 anidulafungin-evolved strains (ANI, ANIFLZ and FinA), 34 developed resistance, of which 33 acquired *FKS1* mutations **(Supplementary Tables 3-5)**. The sole exception (Co_ANI_7F) developed resistance without detectable point mutations or CNVs in known or novel genes compared to its WT strain (**Supplementary Table 5**). This suggested an alternative mechanism not captured by our variant analysis (but see “Drug-induced haploidisation” section).

*FKS1* encodes the catalytic subunit of β-(1,3)-D-glucan synthase, an essential enzyme for the synthesis of β-glucans in the fungal cell wall, and the target for echinocandins^63^. In *Candida* species, echinocandin resistance is typically associated with mutations in two conserved hotspot regions within *FKS* (HS1 and HS2)^27,63,64^. In *C. parapsilosis*, HS1 and HS2 correspond to amino acids 652-660 and 1369-1376, respectively^65^. Among the 29 unique *FKS1* variants identified (**Supplementary Table 6**), most clustered within HS1 (10) or HS2 (7). We also detected three variants in a previously proposed third hotspot region (HS3) in *C. parapsilosis*, spanning amino acids 703-713^66^, including the previously described substitution L703F, previously linked to micafungin resistance, but not to anidulafungin or caspofungin^66^. Our results implicate position 703 in anidulafungin resistance, likely conferring high levels of resistance. Several *FKS1* substitutions recovered here have been reported previously in vitro and/or in clinical isolates (e.g. L703F, W1370R, L1376F; and clinical variants such as S656P, L657W, and R1366H)^66,67,68,69^, reinforcing the clinical relevance of evolution-derived variant catalogues.

Additionally, nine variants occurred outside HS1–HS3. Resistance levels did not differ significantly between strains carrying HS versus non-HS mutations (Wilcoxon rank-sum test, p = 0.121; Supplementary Note 1), indicating that non-hotspot substitutions can also contribute to resistance, albeit with greater variability and likely stronger dependence on genetic background. For example, Cp_FinA_4G carried S1323F and displayed comparatively low resistance among FKS1-mutant strains, consistent with reports that some non-hotspot changes may promote tolerance rather than high-level resistance^65^.**Figure 3B** illustrates the occurrence of novel *FKS1* mutations detected in this study, which represents the first report of unique and shared variants across this species complex and prove the role of these hotspots in *C. metapsilosis* (HS1: amino acids 652-660, HS2: amino acids 1369–1376) and *C. orthopsilosis* (HS1: amino acids 645-653, HS2: amino acids 1362-1369), thus redefining the previously described hotspots for *C. metapsilosis* and *C. orthopsilosis*^70^ (**Figure 3D and F)**.

Beyond *FKS1*, we identified recurrent mutations in genes linked to cell wall stress signalling. *NIK1* (*COS1*) was exclusively mutated in three independent replicates evolved from the *C. metapsilosis* WT BP57 (Cm_ANI_5D, Cm_ANI_5H, and Cm_ANI_5B). *NIK1* encodes a type 3 histidine kinase that negatively regulates the HOG MAPK pathway^71,72^. Although *NIK1* has not been previously directly linked to echinocandin resistance in *Candida*^73^, it plays a role in response to echinocandin-induced cell wall stress, and mutations in this gene conferred resistance to a combined treatment with caspofungin and the metal chelator DTPA^74^. Our finding suggests a potential role of *NIK1* in drug response via modulation of the HOG pathway. Additionally, anidulafungin-evolved Cp_ANI_2C acquired a mutation in *SSK2*, a kinase modulating HOG signaling^75^, underscoring a central involvement of the HOG pathway in echinocandin survival in this species complex. Lastly, we detected a mutation in *RHO1*, a regulatory GTPase that activates β-1,3-glucan synthases, including the *FKS1*-encoded enzyme^76^. This aligns with previous studies linking *RHO1* to echinocandin resistance in *C. parapsilosis*^77,78^. We detected additional low-frequency mutations (**Supplementary Tables 3-5)**, but the concomitant presence of mutations affecting genes previously associated with resistance suggest a passenger or accessory roles.

Altogether, these findings show that anidulafungin resistance in the *C. parapsilosis* complex is largely mediated by a diverse set of *FKS1* mutations, with most (but not all) variants concentrated in conserved hotspot regions. The resulting cross-species mapping of homologous mutations provides a unified framework for interpreting *FKS1* variation in *C. metapsilosis* and *C. orthopsilosis*, where echinocandin resistance mechanisms remain comparatively under-characterised.

### Antifungal exposure frequently induces gene dosage alterations

Protein-altering point mutations are well-established routes to drug resistance and the focus of most previous studies. However, the contribution of gene dosage changes, such as copy number variations (CNVs) and aneuploidies, remains underexplored^41^. We therefore used PerSVade to detect complete and partial aneuploidies, as well as smaller duplications or deletions (**Supplementary Figure 3 and Supplementary Tables 3-5**).

We also attempted to identify Structural Variants (SVs, i.e. inversions, tandem duplications, deletions, insertions and translocations) acquired during adaptation by developing a custom pipeline based on split reads, discordant read pairs and local de novo assembly (see **Methods**). However, after manual inspection of coverage profiles, all candidate SVs classified as “real” by CLOVE were inconsistent with true breakpoints and instead suggested mapping or read-depth artefacts. We therefore did not retain any confident de novo SV calls. Despite this negative result, our simplified, reproducible pipeline for finding changing SVs across microevolutionary timescales may be useful for future work.

In *C. parapsilosis*, we identified 43 CNVs outside aneuploidies, 25 duplications and 18 deletions, affecting 25 and 13 different genes, respectively **(Supplementary Table 8)**. Notably, all duplications were restricted to one strain (Cp_FinA_3D) and involved a single allele. Deletions were mostly concentrated in Cp_CDC317-evolved strains (15/18 deletions) and were mostly associated with anidulafungin treatment (11/15 ANI, 2/15 FinA, and 1 in ANIFLZ, **Supplementary Table 8**). In *C. metapsilosis* and *C. orthopsilosis*, 52 duplications were detected outside aneuploidies. Two *C. orthopsilosis* anidulafungin-evolved strains (Co_ANI_9F and Co_ANI_12A) showed a ∼64 kb triplication in Co_Chr5 spanning 28 genes not previously linked to anidulafungin resistance. Two fluconazole-evolved strains (Co_AinF_11B and Co_AinF_12A) suffered a 46-50 kb duplication in Co_Chr3 that included *MRR1* along with other 17 or 21 genes, respectively. While *MRR1* amplifications have been associated with azole resistance in *C. albicans*^79^, *C. parapsilosis* reported CNVs are restricted to *ERG11* and *CDR1B*^38,48,80^. Importantly, in both *C. orthopsilosis* lineages the duplicated segment included an MRR1 allele that had acquired a resistance-associated point mutation, indicating that CNV and point mutation can act synergistically via increased dosage of a beneficial allele (**Supplementary Table 8**).

Beyond small-scale CNVs, we detected widespread aneuploidies (**Figure 2D**). 39 out of 87 drug-evolved strains carried a segmental or complete aneuploidy, in contrast to no aneuploidy in WT and YPD-evolved strains (**Supplementary Tables 3-5**). Aneuploidy was frequently unstable and extra chromosomal material was lost in 70% (9/13) of switch conditions, consistent with high karyotypic plasticity^36,81,82^. Whole-chromosome or partial trisomy dominated, with only two identified tetrasomies (but see “Drug-induced haploidisation” section) and one segmental deletion in Cm_Chr7.

Many segmental aneuploidies involved a single chromosomal arm (**Supplementary Figure 3A**), consistent with frequent centromere rearrangements in the *C. parapsilosis* complex^83^. Specific homologous chromosomal arms showed recurrent duplication under antifungal stress across different species, most prominently the long arm of Cp_Chr1 and the corresponding homologous regions in other species (**Supplementary Figure 3B**). Similar recurrent aneuploidies have been reported in *C. albicans* isolates tolerant to fluconazole^84^ and in caspofungin-tolerant *C. parapsilosis*^36^.

Aneuploidies were almost twice as frequent under fluconazole exposure as compared to anidulafungin (50% of FLZ and AinF conditions vs. 27% of ANI and FinA conditions). More than half of fluconazole-exposed aneuploid strains (10/19) had dosage changes affecting the region containing *ERG11, TAC1* and *CDR1B*, a region never affected under anidulafungin exposure (**Figure 4**). This pattern is reminiscent of azole-associated isochromosome 5L formation in *C. albicans*^85^, and could explain Co_AinF_7F resistance (**Supplementary Table 5**). Other fluconazole-exposed strains showed aneuploidies recurrently affecting *MDR1B* (7/19) and *MRR1* (5/19), consistent with previous observations^79^. Moreover, aneuploid segments often affected alleles carrying newly acquired MRR1 or ERG11 point mutations, again suggesting selection for increased dosage of adaptive alleles. Overall, 89% of fluconazole-exposed strains that became aneuploid had gene dosage alterations affecting one of the key resistance genes (*MRR1*, *MDR1B, ERG11, TAC1* or *CDR1B*).

**Figure 4.**
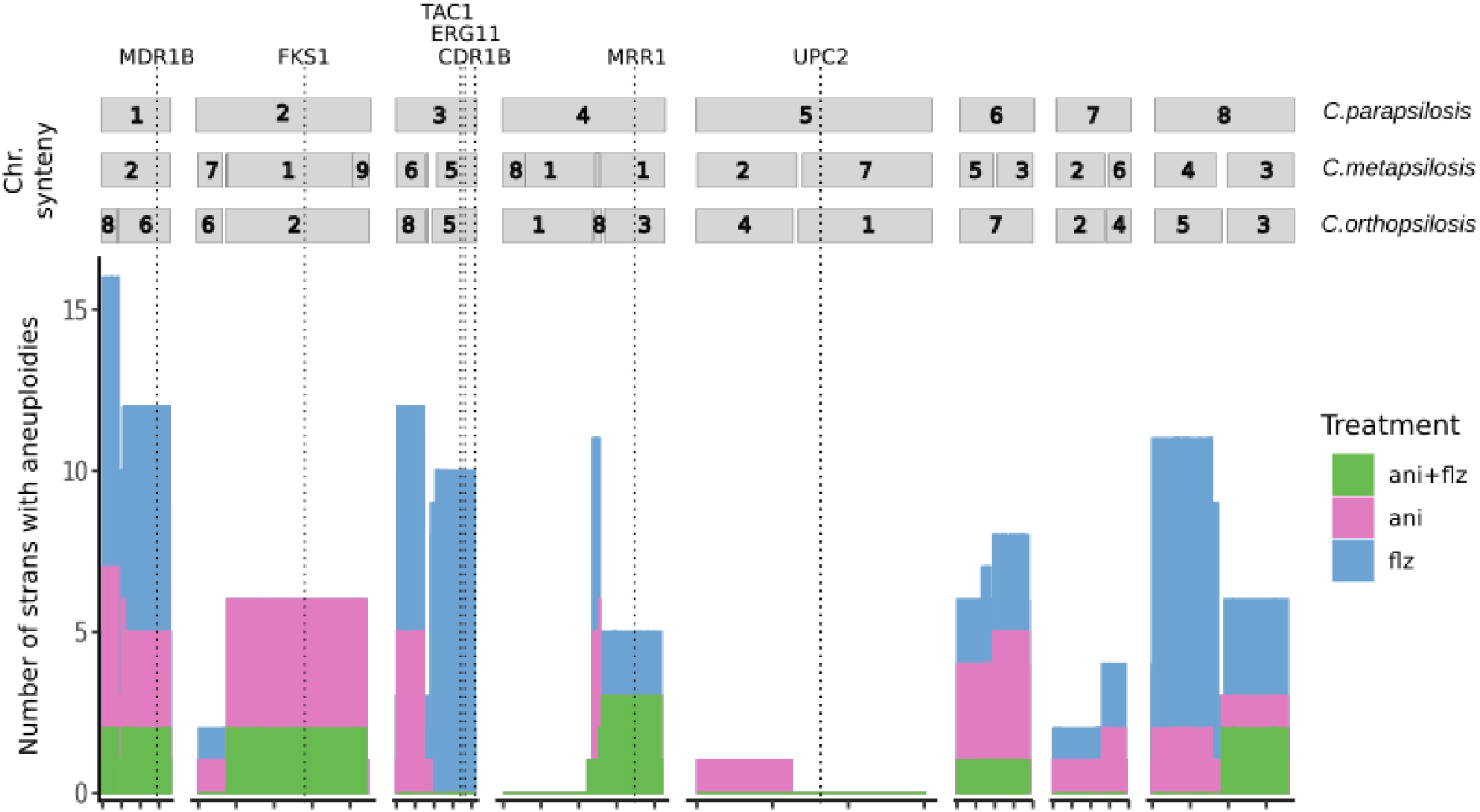
Aneuploidies recurrently affect genes related to drug resistance. Frequency of aneuploidies in evolved strains of the *Candida parapsilosis* complex. Synteny between the three species and the localization of key resistance genes is represented above.

Under anidulafungin treatment we observed duplications of Cp_Chr2 and its homologous region in Co_Chr2, encoding *FKS1* (**Figure 4**), linking these aneuploidies with anidulafungin adaptation. Similarly, we detected a partial aneuploidy in the homologous region of Cp_Chr5 (Co_ANI_10E), consistent with its identification in caspofungin-tolerant *C. parapsilosis* isolates^36^.

Hence, CNVs and aneuploidies are a frequent outcome of antifungal exposure in the *C. parapsilosis* complex. The recurrence of dosage changes affecting homologous regions across species and treatments argues against purely stochastic stress-induced imbalance^34^ and instead supports a substantial adaptive contribution through increased dosage of key resistance loci. These changes were often reversible once drug pressure was removed, consistent with fitness costs in drug-free environments.

### Gene conversion via LOH favours adaptive alleles in heterozygous backgrounds

Loss of heterozygosity has been proposed in *C. albicans* to expedite drug resistance by favoring gene conversion towards mutant alleles in genes such as *TAC1*^86^, *UPC2*^87^, *MRR1*^49^ or *FKS1*^88,89^. Additionally, in highly-heterozygous hybrid strains, LOH can resolve negative allelic epistatic interactions^39^. We therefore investigated the occurrence of LOH during drug exposure in *C. orthopsilosis* and *C. metapsilosis* hybrids, using the SNP-density approach implemented in JLOH^90^.

Across evolved hybrids, newly acquired LOH covered on average 0.4% of the genome, with rare cases reaching ∼15% (excluding haploidised strains; (see Drug-induced haploidisation section). The extent of acquired LOH was not different between species nor between drug-evolved and YPD-evolved strains (**Supplementary Note 1**), consistent with studies in *C. albicans* showing no induction of LOH by fluconazole^91^.

Despite the overall limited LOH, some key genes were recurrent targets (**Figure 2E**). Strikingly, mutations acquired in *MRR1* or in *FKS1* were frequently rendered homozygous within an LOH block (17/26 and 7/10, respectively, **Supplementary Table 7**), supporting selection of gene conversion towards the resistant allele. In contrast, compensatory *MRR1* mutations arising during switch conditions (FinA) were usually (3/4) heterozygous and never part of a newly formed LOH. Together, these results suggest that antifungal exposure does not broadly increase short-range LOH, but that selection favours LOH events that fix previously acquired resistance mutations, as proposed in other *Candida* species^92,93^.

### Drug-induced haploidisation facilitates antifungal adaptation

Unexpectedly, LOH profiling revealed several strains with apparent genome-wide LOH (Co_ANIFLZ_10E, Co_ANI_10G, Co_AinF_10G, Co_FLZ_10A, Co_FinA_10A) (**Figure 5A-B, Supplementary Figure 4**). We hypothesised that these patterns reflected haploidisation rather than extensive LOH tracts (**Figure 5A**). This interpretation was further supported by their low fitness phenotype and the high number of apparent tetrasomies, which are more parsimoniously explained as disomies on an haploid background (**Supplementary Table 5**). Similar phenotypes were also observed in some non-hybrid strains not included in the LOH analysis (Co_ANI_7F, Co_FLZ_12A, Co_FLZ_12G, and Co_ANI_12C).

**Figure 5.**
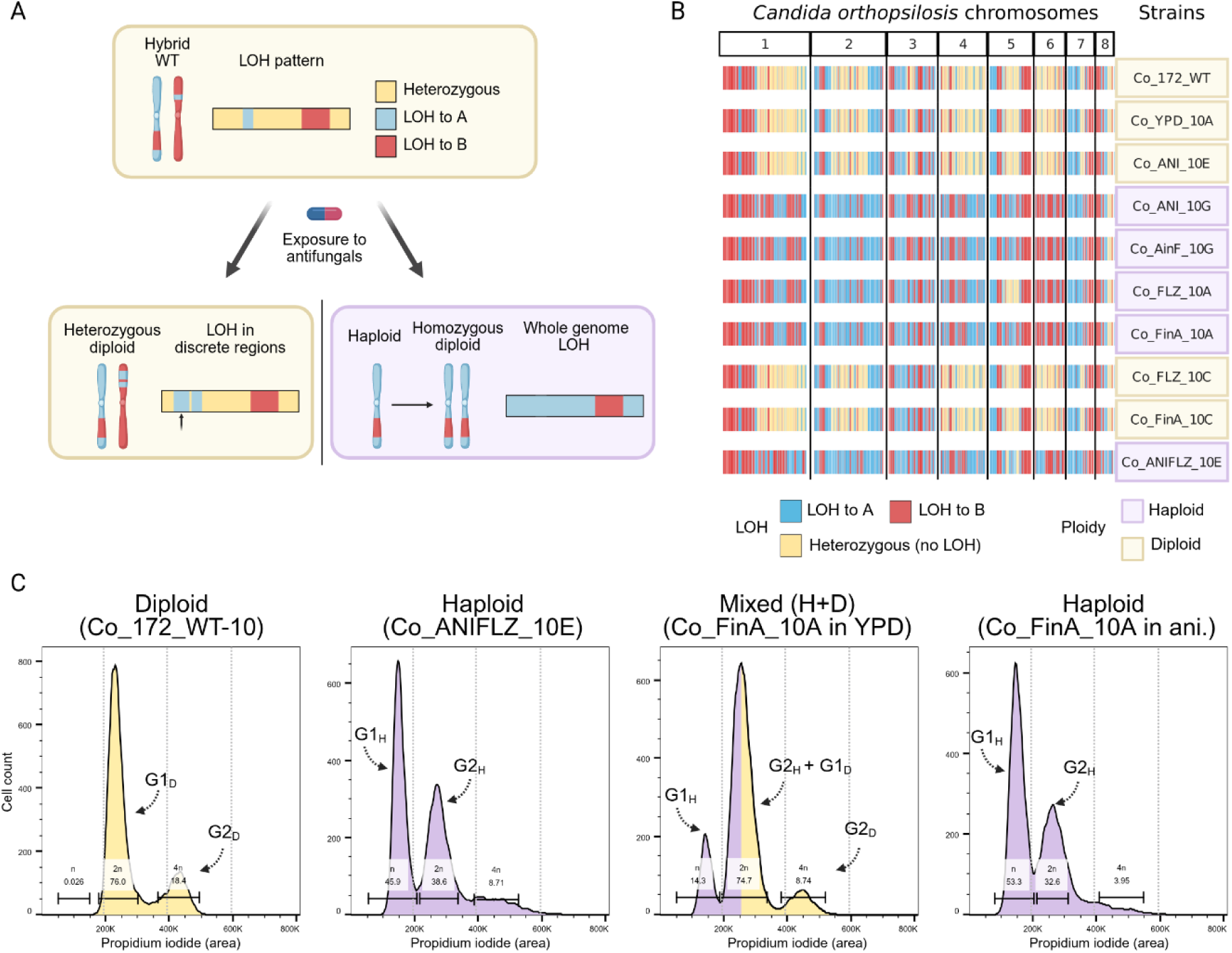
Whole-genome LOH as a consequence of haploidisation. **(A)** Schematic representation of the evolutionary trajectories and LOH patterns of a heterozygous hybrid towards short-range LOH acquisition (left) or haploidisation and later whole-genome duplication (right). **(B)** LOH patterns observed in evolved isolates of *C. orthopsilosis* 172. (**C)** Propidium iodide area (PI-A) for living cells of *C. orthopsilosis* 172. The area is coloured according to the assigned ploidy (diploid: yellow, haploid: lilac, mixed: both). For each peak, cell cycle phase (G1, G2) is indicated for haploid (H) or diploid (D) cells.

We validated ploidy by flow cytometry^94^, where haploid cells are expected to show G1(n) and G2/M(2n) peaks at half the DNA content of control diploids (**Figure 5C and Supplementary Figure 5A**). For most suspected haploids, the G1 peak was ∼50% of the corresponding diploid WT, whereas the G2/M peak aligned with the WT G1 peak (**Figure 5C; Supplementary Table 9**), confirming haploidisation. Notably when propagated without drugs, some haploid strains exhibited predominantly diploid (Co_AinF_10G and Co_ANI_12C) or mixed haploid/diploid (Co_FinA_10A and Co_ANI_7F) profiles, indicating that haploidisation is reversible. This spontaneous rediploidization could explain why some suspected haploids showed a diploid profile. In contrast, growth under drug pressure (fluconazole 128 µg/ml and/or anidulafungin 2 µg/ml), maintained predominantly haploid populations (**Figure 5C and Supplementary Table 9**). This finding indicates that haplody is a transient, drug-selected state that is disfavored in drug-free conditions, consistent with instability and high fitness costs^95^.

Interestingly, one fluconazole-evolved haploid strain (Co_FLZ_10A) harbored a heterozygous aneuploidy in Co_Chr5 including *ERG11*-*TAC1*-*CDR1B* (**Figure 5B, Supplementary table 5**), indicating that the two original copies of the chromosome were retained despite haploidisation and reinforcing the relevance of this chromosome in fluconazole resistance.

Haploid lineages arose across different clades and drug treatments, despite the severe fitness costs, indicating convergent selection for ploidy reduction as a survival strategy **(Figure 2F)**. This is exemplified by strain Co_ANI_7F, the sole anidulafungin-evolved lineage exhibiting cross-resistance to fluconazole, which is haploid and lacks other identifiable resistance-conferring mutations (**Supplementary Table 5)**. Similarly, strains displaying increased tolerance to anidulafungin (Co_ANI_10E, Co_ANI_10G, Co_ANIFLZ_10E, and Co_FinA_10A) were devoid of canonical FKS1 mutations but had uniformly undergone haploidisation (**Supplementary Table 5, Supplementary Figure 2**).

While there are previous reports of fitness-compromised haploid strains in *C. albicans*^96^, and azole-resistant haploids in *C. tropicalis*^95^, the selective advantage of ploidy reduction remains unclear.^95^ We propose three non-mutually exclusive explanations. First, haploidisation provides a rapid path to homozygosity, potentially unmasking recessive resistance alleles. Second, it may represent a persistence state analogous to slow-growing, metabolically reduced phenotypes (e.g. small colony variants)^97–99^, consistent with low-fitness observed here and transcriptomic signatures reported for azole-resistant haploids in *C. tropicalis*^95^. Third, as described in other yeasts^100^, haploid cells may activate distinct transcriptional programs that would be advantageous under drug stress. Detecting such events solely from genome data in highly homozygous diploid species like *C. parapsilosis* would be very challenging. This may explain why this mechanism has remained undiscovered until now. Together, our results identify drug-induced, reversible haploidisation as a previously unidentified adaptive mechanism in the *C. parapsilosis* species complex.

### Pervasive fitness costs shape resistance stability

Fitness costs strongly impact survival and spread of resistant strains, by reducing their ability to compete with WT populations in drug-free environments^44,101^. Fluconazole-resistant outbreaks have led to suggestions that certain *C. parapsilosis* resistant strains may carry little or no fitness costs^27^. However, this remains unproven. Our experimental design allows pairing each evolved lineage with its drug-naïve WT ancestor, enabling direct quantification of fitness consequences attributable to resistance acquisition. We measured growth in rich medium (YPD) and defined relative fitness (AUC_R_) as the ratio of AUC of each evolved strain over the corresponding WT, where values close to 1 indicate fitness similar to the WT strain and values above or below 1 indicate increased or decreased fitness, respectively.

Our results show compelling evidence that significant fitness reduction is a widespread trade-off of drug adaptation in all species of this complex (**Figure 6A**). Control YPD-evolved strains showed no fitness change relative to WT (two-sided Wilcoxon signed-rank test, ns), whereas all drug-evolved strains exhibited significant fitness reduction (two-sided Wilcoxon signed-rank test, FDR-adjusted p ≤ 0.017; **Supplementary Note 1**), as indicated by arrows in **Figure 6A**. FLZ strains experienced the most pronounced fitness loss in monotherapy, with an average reduction of ∼70%, while ANI strains showed a milder cost (∼25%). Drug switch experiments further supported this trend. ANI strains subsequently exposed to fluconazole (AinF) experienced a significant reduction in fitness (**Figure 6A, C**), while FLZ strains exposed to anidulafungin (FinA) showed a surprising recovery in fitness (**Figure 6A, D**), likely due to the appearance of compensatory mutations. Strains in the combined treatment (ANIFLZ) suffered the greatest fitness costs (∼90%), underscoring the compounding stress of simultaneous resistance and explaining the low survival rate under these conditions.

**Figure 6.**
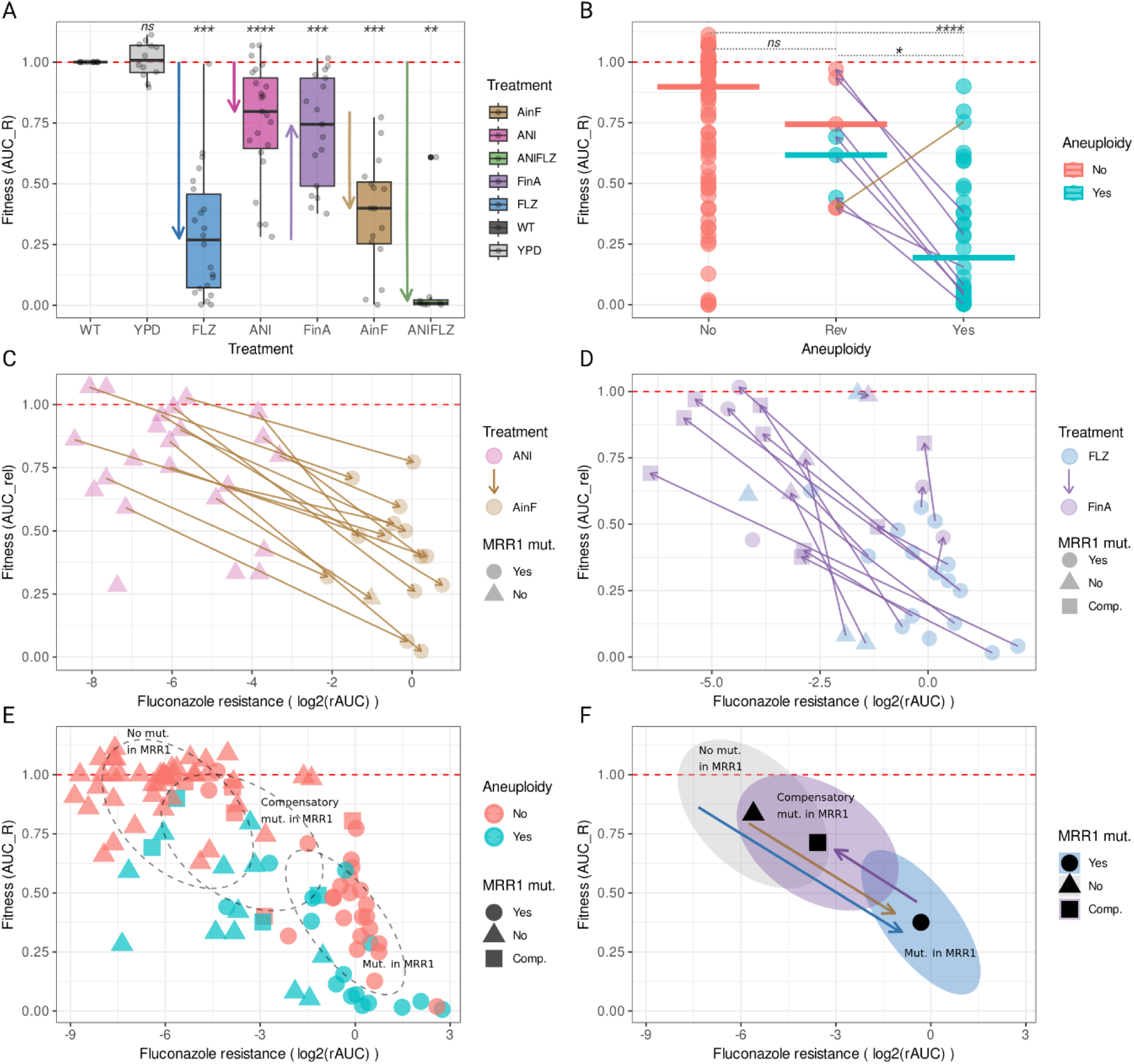
Fitness costs are associated with drug resistance and are a critical factor for its stability. Fitness (AUC_R_) is the ratio between the AUC in YPD of the evolved strain and the AUC in YPD of their WT strain. The red dotted line indicates the fitness of the WT as a reference, so if it is above, the fitness is increasing, and if it is below, it means that they have decreased their fitness. In panels (**C-F**), resistance to fluconazole was measured as log2 of the resistanceAUC (rAUC) and point shape indicates the presence of missense mutations (Yes), stop mutations or compensatory missense mutations (Comp.) or the absence of mutation (No) in *MRR1*. (A) Changes in fitness of strains evolved in the different treatment conditions. Arrows emphasise the median fitness changes in each treatment between the parental and evolved strains. In boxplots, the central line indicates the median, the box spans the interquartile range (Q1 to Q3), and whiskers extend to the smallest and largest values within 1.5×IQR. The points represent the median between 4 biological replicates per strain. (**B**) Effect of aneuploidies in relative fitness (AUC_R_). The x axis groups evolved strains without aneuploidies (No), with aneuploidies (Yes), and which lost some or all of their aneuploidies in the switch conditions (Rev, connected with arrows to the strain from which it originated). Colours indicate whether the strain has any aneuploidies. (**C**) Fitness (AUC_R_) of strains evolved in anidulafungin (ANI) and subsequently in fluconazole (AinF) conditions versus resistance to fluconazole (log2(rAUC)). Arrows connect paired ANI and its switch evolved AinF strains. (**D**) Fitness (AUC_R_) of strains evolved in fluconazole (FLZ) and subsequently in anidulafungin (FinA) conditions versus resistance to fluconazole (log2(rAUC)). Arrows connect paired FLZ and its switch evolved FinA strains. (**E**) Fitness cost associated with FLZ resistance and its origins. Dotted ellipses group 50% of samples according to the presence and type of *MRR1* mutations. (F) Dynamics of fitness cost in the development of fluconazole resistance and later compensatory mutations in switch conditions. Coloured ellipses group 50% samples according to the presence and type of *MRR1* mutations. Arrows show the overall trend in fitness and resistance from WT to FLZ (blue), ANI to AinF (ocher) and FLZ to FinA (purple). Asterisks indicate significance levels based on adjusted p-values (corresponding statistical tests for **A, B** see Methods): * indicates p < 0.05; ** indicates p < 0.01; *** indicates p < 0.001; and **** indicates p < 0.0001.

Aneuploidy was a major determinant of fitness (Kruskal-Wallis test p = 4.79×10^-10^; **Figure 6B**), with aneuploid strains having significantly lower fitness compared to both euploid strains (Dunn-FDR p.adj = 1.71×10^-10^) and strains that had reverted to euploidy (Dunn-FDR p.adj = 0.031), while no significant difference was observed between reverted and euploid strains (Dunn-FDR p.adj = 0.22), regardless of treatment conditions (**Supplementary Note 1**). This likely reflects the burden imposed by the increased dosage in hundreds of genes^102,103^. In contrast, anidulafungin resistance driven solely by *FKS1* mutations (n=18), was associated with comparatively milder costs retaining ∼88% of their WT fitness (**Figure 6A**). Among anidulafungin-evolved lineages, the greatest costs were observed in multiresistant strains combining *FKS1* and *MRR1* mutations and/or aneuploidies (**Supplementary Figure 6C)**.

For fluconazole, resistance levels correlated with fitness costs (**Figure 6E**). Strains with *MRR1* mutations were consistently more resistant to fluconazole, but had the most severe fitness loss (average loss ∼68%). Tracking individual strains during drug switch treatments revealed that resistance gained through *MRR1* mutations invariably came at the expense of fitness, with significant reductions seen in every case (**Figure 6C**, two-sided paired t-test, p = 3.17 × 10^-7^; **Supplementary Note 1**)).

We hypothesised that resistance-associated fitness trade-offs could be relevant determinants of resistance stability. To test this, we analyzed drug-switching regimes, which mimic changes in therapy following treatment failure. We observed two contrasting outcomes. Anidulafungin resistance remained largely stable after switching to fluconazole (AinF, **Supplementary Figure 1B, Supplementary Figure 6A**). In contrast, fluconazole resistance decreased significantly after switching to anidulafungin treatment (FinA, **Supplementary Figure 1A, Figure 6D**). Supporting our departing hypothesis, this asymmetry parallels the fitness costs of the corresponding resistance mechanisms: high-cost fluconazole resistance becomes disfavoured when fluconazole pressure is removed, promoting compensatory changes that restore fitness and concomitantly reduce resistance.

To understand the genetic basis of resistance loss, we inspected mutations appearing during FinA switch conditions. Notably, the appearance of additional *MRR1* mutations correlated with improved fitness and reduced fluconazole resistance (**Figure 6D),** suggesting a compensatory role. Resistance loss was often partial, with FinA derivatives typically showing intermediate between FLZ and WT(**Figure 6E).** This intermediate pattern is consistent with compensatory mutations remaining heterozygous **(Supplementary Table 7)**, whereas resistance-associated mutations were usually homozygous (see above), suggesting that complete homozygosity of compensatory mutations may be detrimental. A comparable trajectory was observed in clinical isolates of *Clavispora (Candida*) *lusitaniae*^104^, where resistance-conferring gain-of-function mutations were followed by secondary loss-of-function mutations that reduced resistance.

Fitness restoration in switch treatments also occurred via aneuploidy reversion, with strains losing aneuploid segments significantly improving fitness (Kruskal-Wallis test followed by Dunn’s post-hoc with FDR correction, p. adj = 0.033) (**Figure 6B**). This was most pronounced in FinA, where seven aneuploidies were reverted, five of which involved the chromosome containing *ERG11* and *TAC1*, consistently linking the loss of resistance to the increase of fitness. By contrast, only one aneuploidy reversion was observed in AinF (Co_ANI_9H), and this lineage simultaneously acquired an MRR1 mutation, potentially offsetting any fitness benefit. Together with the rediploidization process described above, these results underscore the important role of trade-offs in shaping drug resistance stability: when drug pressure is removed, fitness-restoring reversion and compensatory evolution are favoured, leading to reduced (often partial) resistance.

## Conclusions

By integrating experimental evolution, genome sequencing, and phenotyping across the phylogenetic breadth of the *Candida parapsilosis* species complex, we show a conserved repertoire of antifungal adaptation strategies that is predictable in its molecular targets but highly plastic in its genomic execution. Across species, backgrounds and isolation niches, resistance repeatedly converged on a shared set of canonical genes, being *MRR1* and *FKS1* the dominating targets of azole and echinocandin adaptation, respectively. Although protein-altering point mutations, generally arising within hotspot regions were the most common adaptation mechanism, they were frequently reinforced or substituted by plastic gene-dosage changes (CNVs and aneuploidy) and, in hybrids, by LOH that drives resistance alleles to homozygosity. Our results provide an expanded catalogue of resistance-conferring mutations and a unified framework to interpret resistance evolution in *C. parapsilosis*, and the comparatively undercharacterised *C. metapsilosis* and *C. orthopsilosis*. Moreover, they also caution that supra-MIC survival and tolerance may enable persistence even when stable resistance is not detected by routine testing.

Consistent with higher clinical prevalence, *C. parapsilosis* exhibited higher survival rates as compared to *C. orthopsilosis* and *C. metapsilosis*, although hybrid strains of *C. orthopsilosis* showed a better adaptability profile compared to non-hybrid strains. This provides first evidence of adaptive advantage of hybrid strains in clinical settings, something previously anticipated but not demonstrated^10,11^. Compared to a similar experiment in *N. glabratus*^43^, our results show a more limited capacity of evolving cross– and multidrug-resistance in this species complex, consistent with lower rates of multidrug resistance among clinical isolates^105^. Notably, combination therapy resulted in the highest extinction rates and effectively prevented the emergence of multidrug resistance, supporting its superior suitability for clinical practice.

Most notably, we uncover drug-selected, reversible haploidisation as a novel adaptive route to antifungal survival. Its recurrence across clades and treatments, its association with strong fitness defects, and its reversion in drug-free conditions together indicate that ploidy reduction can act as a transient stress-adaptive state. This expands the catalogue of resistance and tolerance-associated genomic strategies beyond point mutations and copy-number changes, and highlights that clinically relevant adaptive states may be missed if surveillance focuses solely on canonical SNP markers in diploid genomes.

Finally, our data show that resistance, especially high-level fluconazole resistance, is commonly accompanied by substantial fitness costs, promoting compensatory evolution once selection is removed. This predicted evolutionary outcome provides interesting new insights for therapy design^106,107^, suggesting that drug-switch strategies^108^ may be particularly effective in mitigating azole resistance in these species. This also raises the question as to whether the frequently observed *MRR1*-mediated resistance could underlie therapy failures and later be reverted or not detected due to the high fitness costs in the absence of drug exposure. Together, our work provides a cross-species, genome-wide reference for resistance surveillance and a conceptual framework for developing novel therapeutic approaches in this pathogen complex.

## Methods

### Strains and growth conditions

We selected twelve strains of the *C. parapsilosis* complex representing a diversity of genetic, geographical and niche origins (**Supplementary Table 1**). For each strain, we isolated a single colony by streaking them in agar plates from 20% glycerol stocks kept at –80°C. Unless indicated otherwise, we grew strains in YPD medium (1% yeast extract, 2% bactopeptone, 2% glucose) with 2% agar for solid cultures, shaking at 200 rpm liquid cultures, and incubated at 30°C for 24 hours.

### *In vitro* evolution experiment

In vitro evolution experiments were conducted as previously described^43^ with minor modifications. In brief, we evolved strains in 96 deep-well plates (Sarstedt), each well containing the required medium and a 3 mm glass bead (VWR) for agitation. We used a sandwich cover (Enzyscreen BV) to ensure proper aeration while minimizing evaporation. We cultured each of the 12 parental (wild type, WT) strains in a different column of the plate, assigning each WT the number of the corresponding plate column. For each WT we distributed four biological replicates in four rows of the corresponding column, and assigned the corresponding row letter to that replicate name. In each row we included empty controls (non-inoculated medium) and experimental wells, alternating them in a checkerboard pattern to prevent cross-contamination. At each passage, we assessed signs of potential contamination in empty control wells. At the start of the evolution experiment, passage 0 (P0), we inoculated single WT colonies in YPD for 48 hours at 30°C and 200 rpm. From this P0 plate, we transferred 50 µl of culture into treatment plates containing 450 µl of YPD supplemented with either anidulafungin (ANI treatment), fluconazole (FLZ treatment), the combination of both drugs (ANIFLZ treatment), or a YPD control (YPD treatment). For each condition, in each passage, we transferred 50 µl of culture into 450 µl of fresh medium with the corresponding treatment every 48 hours – three times per week – for a total of 18 passages. We doubled drug concentrations in alternate passages, starting at 0.016 µg/ml and increasing to 4.096 µg/ml for anidulafungin and from 4 µg/ml to 192 µg/ml for fluconazole. The same concentrations were used for the ANIFLZ treatment. Prior to each drug concentration increase glycerol stocks were prepared to allow for recovery of a population at a desired time point.

At the end of passage 18 (P18), we plated each sample on YPD medium containing the final drug concentrations. We randomly selected a single colony from each plate, grew it in liquid YPD supplemented with the same drug concentration, and stored in glycerol until further use. We extracted genomic DNA from these evolved isolates for sequencing, and used them as parental strains for the second round of evolution.

In the second round of evolution we switched drug treatments so that strains previously evolved in fluconazole were evolved in anidulafungin (FinA treatment) and vice versa (AinF treatment). Otherwise, we used the same protocol and drug concentrations as described above.

We named evolved strains with unique alphanumeric identifiers indicating the treatments (YPD, ANI, FLZ, AinF, FinA, ANIFLZ), followed by their corresponding plate coordinates. These coordinates, in turn, indicate the WT strain as column number (1-12) (**Supplementary Table 1**) and biological replicates as rows (B, D, F, H for even rows and A, C, E, G for odd rows). For clarity, in the text we added the species initials before the code. For example, Cp_FLZ_1H refers to a *C. parapsilosis* (Cp) strain, evolved in fluconazole (FLZ) from the WT strain GA1 (assigned to number 1), and is the fourth replicate (H) of this condition. In addition, each strain was assigned a unique numerical code (1-288) that is omitted from the text for simplicity.

### Fitness measurement and antifungal susceptibility testing

We used Q-PHAST^45^ to measure fitness in YPD medium (control) and in YPD supplemented with 11 concentrations (2-fold increased) of fluconazole (ranging from 0.25 to 256 ug/ml) or anidulafungin (from 0.016 to 16 µg/ml). In brief, we grew four biological replicates for each evolved strain, and distributed them on a 96 deep-well Master-Plate (MP) according to the Q-PHAST replicate distribution^45^, which minimizes position-dependent biases. We placed evolved strains and its corresponding WT strain in the same plate for optimal comparison. After 24 hours incubation, when cultures reached saturation, we diluted 3 µl of each sample in 200 µl of water in a Dilution-Plate (DP), from which we placed 96 spots of 3 µl on agar plates containing the drugs, referred to as Experimental-Plates (EPs). EPs were incubated at 30°C on top of scanners that captured images every 15 minutes for 24 hours. We analyzed these images, along with strain information introduced in the Plate-Layout (PL), using the Q-PHAST software (v1), obtaining growth curves for each spot. We used the area under the curve (AUC) as a proxy for fitness for each spot at each tested condition.

For each evolved strain we divided its AUC in the control (YPD medium) by the AUC of the corresponding WT strain in the same control, obtaining the AUC relative to its WT (AUC_R_), which measures the fitness cost of resistance for each strain. In addition, the growth of each strain at each drug concentration was calculated relative to its fitness in the control medium, and a curve representing the relative fitness of this strain across all drug concentrations was constructed^43,45^. From this curve of relative fitness, the area under the curve, called resistance AUC (rAUC), and minimum inhibitory concentration 50 (MIC_50_) were used as susceptibility measures, for which we calculated the median (M) and median absolute deviation (MAD) from the four replicates. Resistance AUC (rAUC) susceptibility values correct for genetic background differences and specifically inform on variations on fitness or drug susceptibility resulting from mutations acquired during the evolution experiment^43^. rAUC values near 1 indicate similar fitness across all tested concentrations, hence low drug susceptibility, while lower values reflect higher susceptibility. Unlike MIC_50_, which captures a single inhibition point, rAUC provides a broader quantitative measure of growth across the entire concentration range^45^. We additionally computed the relative MIC_50_ (MIC_R) by dividing the MIC_50_ of the evolved strain to the MIC_50_ of its corresponding WT. Evolved strains with a MIC_R value equal to or greater than 4 (indicating at least a 4-fold decrease in susceptibility) were considered as resistant (R), whereas those with a lower value were deemed sensitive (S), given that all WTs were drug sensitive. Susceptibility assessment was not possible for 8 out of 111 strains due to insufficient growth within 24h.

To re-evaluate susceptibility in anidulafungin-evolved survivors lacking resistance mutations, 96-well broth microdilution assays were performed in YPD medium supplemented with twofold serial dilutions of anidulafungin. Wells were inoculated with 1.25×10^5^ fresh cells and incubated at 30 °C. MIC_50_ was determined after 24 hours. To assess fungicidal activity, 3 µl aliquots from each well were spotted onto drug-free YPD agar plates. Liquid plates were incubated for an additional 24 hours (48 hours total) to determine the 48-h MIC_50_ and to quantify Supra-MIC Growth (SMG), defined as the average growth at concentrations exceeding the 24-h MIC_50_. Resistance ratios (RR) were calculated by normalizing the MIC_50_ of evolved strains to their respective parental WT (MIC_evolved_/ MIC_WT_). Finally, 100 µl aliquots from wells corresponding to 4 µg/ml 48 hours of incubation plate were plated onto drug-free YPD agar plates to quantify viable Colony Forming Units (CFUs) and evaluate cell viability under conditions mimicking the evolution experiment. The experimental design is illustrated in **Supplementary Figure 2A**.

### Determination of ploidy by flow cytometry

To assess changes in ploidy, defined as the number of sets of homologous chromosomes within a cell, we quantified the cellular DNA content using flow cytometry following a previously described protocol with minor modifications^94^. Briefly, test and control strains were streaked on solid medium and incubated for 72 hours at 30°C. Cells were then cultured overnight in liquid medium at 30°C 200 RPMs with or without antifungal drugs, as described. Cultures were adjusted to optical density (OD₆₀₀) of 0.8, centrifuged, and washed in 50 mM sodium citrate buffer solution. Cells were fixed in 70% (v/v) ethanol for at least 1 hour at room temperature. Following fixation, cells were washed and treated with RNase A (0.5 mg/ml in 50 mM sodium citrate containing 40% (v/v) glycerol) for 2 hours. DNA was stained overnight in the dark with 7.5 µl of 1 mg/ml propidium iodide (PI) solution in water and 5 µl of 50 mM sodium citrate. Prior to flow cytometry analysis, cells were sonicated using a Bioruptor (Diagenode) sonicator for 20 cycles of 10 seconds ON /30 seconds OFF at low power to minimize cell clumping while preserving cell integrity. Cells were diluted 1:10 in 25 µg/ml PI in 50 mM sodium citrate immediately before analysis.

Samples were analyzed using a Gallios multi-color flow cytometer instrument (Beckman Coulter, Inc, Fullerton, CA) set up with the 3-lasers 10 colors standard configuration. Excitation was done using a blue (488nm) laser. Forward scatter (FS), side scatter (SS), red (620/30nm) fluorescence emitted by propidium Iodide (PI) was collected. At least 20,000 events per sample were recorded at a flow rate of ∼300 cells per second. Data were analyzed using FlowJo v10 software^94^. LMD files were imported into a single worksheet, and a standardized gating strategy was applied across all samples (**Supplementary Figure 5**). To exclude cell debris and doublets, a gate was defined on a PI-Area vs. PI-Height plot (Comp-FL3-A:: FL3 INT LIN vs. Comp-FL3-H:: FL3 INT LIN), selecting single cells aligned along the vertical axis. Gates were applied uniformly across all samples, with minor adjustments to ensure optimal fitting. Additional scatter plots (FSC-H vs. SSC-H and PI-A vs. FSC-H) were used to assess cell size and complexity. For each gated population, PI-A histograms were generated to evaluate DNA content and infer cell cycle phases (G1 and G2/M). The G1 fluorescence peak was manually defined using the WT diploid control strains (2n). Peaks at half (n) or double (4n) the WT G1 intensity were defined, with minor adjustments for aneuploidies. Strains with G1 peak at 2n were classified as diploid and strains showing a peak at n (>1,000 cells) were classified as haploid. Populations with peaks at n, 2n, and 4n (>1,000 cells) were classified as a mixture of haploid and diploid cells.

Examples of the gating and analysis strategy are shown in **Supplementary Figure 5.** Frequencies and single-cell counts for each ploidy state (n, 2n, and 4n) are provided in **Supplementary Table 9**.

### DNA extraction, library preparation and sequencing

We extracted genomic DNA from yeast cultures with the YeaStar Genomic DNA Kit (Zymo Research) following manufacturer’s instructions. Briefly, cultures were grown overnight in 15 ml YPD on an orbital shaker at 200 rpm and 30°C. We harvested cells by centrifugation 2ml of culture at 500g for 2 minutes. Pellets were resuspended by vortexing with 120 µl YD Digestion Buffer and 5 µl R-Zymolyase and incubated at 37°C for 60 minutes. Then 120 µl of YD Lysis Buffer was added, followed by vigorous vortexing and 250 µl of chloroform. After thorough mixing, we centrifuged samples at 10,000 rpm for 2 minutes and transferred the supernatant to a Zymo-spin III column. We then centrifuged the columns at 10,000 rpm for 1 minute. Subsequently, we washed the samples twice with 300 µl DNA Wash Buffer and centrifuged them at 10,000 rpm for 1 minute each time. Finally, we transferred the Zymo-spin III columns to new tubes, added 60 µl of TE and, after 5 minutes incubation, we centrifuged them at 10,000 rpm for 30 seconds to elute the genomic DNA.

Genomic DNA libraries for whole genome sequencing were prepared at the IRB Barcelona Functional Genomics Core Facility. At least 650 ng of genomic DNA dissolved in a final volume of 50 µl TE buffer, whenever available, or at least 100 ng were sheared with a Bioruptor sonicator (Diagenode) using the following settings: temperature 4-10 °C; intensity: high; cycles: 3; cycle time: 5 minutes; cycle program: 30 seconds pulse and 30 seconds rest time. Additional one-minute fragmentation cycles (30 seconds on/off) were performed when necessary to ensure homogeneous fragmentations. After each sonication cycle, samples were centrifuged at 4 °C and the water tank was refilled with pre-cooled water. DNA fragmentation was quality controlled using the Bioanalyzer 2100 and its DNA High Sensitivity chip (Agilent) and quantified using the Qubit fluorometer and its dsDNA HS assay (Invitrogen). NGS libraries were prepared from 250 ng of fragmented genomic DNA, whenever available, or from 100 ng using the NEBNext Ultra II DNA library prep kit for Illumina (New England Biolabs). Adaptor-ligated DNA were size-selected using the provider-recommended settings to obtain an insert size distribution of 300-400 bp. After purification, libraries were amplified through five or seven amplification samples, depending on the input material used, using the NEBNext multiplex oligos for Illumina (New England Biolabs). The final libraries were quantified on Qubit and quality controlled in the Bioanalyzer. An equimolar pool was prepared with the number of samples included in each subproject, and submitted for sequencing at the Centre Nacional d’Anàlisi Genòmica (CNAG). A final quality control by qPCR was performed by the sequencing provider and libraries were sequenced on a NovaSeq6000 (Illumina) using the paired-end 150 nt strategy.

### Read mapping and variant calling

We used perSVade (v. 1.02.6)^109^, a pipeline including modules for sequencing read trimming, quality assessment, alignment, variant and copy number variant (CNV) calling, as well as for functional annotation of the identified variants. For adaptor removal and read trimming we used FastQC (v. 0.11.9)^110^ and Trimmomatic (v. 0.38)^111^ with default parameters, included in the trim_reads_and_QC module of perSVade. This was followed by alignment of the trimmed reads with BWA MEM (v. 0.7.17) to the reference genomes of *C. parapsilosis* CDC317 (ASM18276v2, GCA_000182765.2), *C. orthopsilosis* Co90-125 (ASM31587v1, GCA_000315875.1^112^) or *C. metapsilosis* BP57 (GCA_017655625.1^39^), using the align_reads module of perSVade. The *C. parapsilosis* reference is identical to the one available at the Candida Genome Database (CGD, v. s01-m06-r07). The correspondence of chromosome names between different databases and the names used in this paper is provided in **Supplementary Table 10**. Next, using the module call_small_variants from perSVade we performed variant calling on the aligned reads and annotated the function of small variants (SNPs and small indels). The results integrate the output from three different variant callers: BCFtools (v. 1.9)^113^, GATK Haplotype Caller (v. 4.1.2)^114^, and Freebayes (v. 1.3.1)^115^. For variant calling, we set the ‘--ploidy 2’ option, which is the canonical ploidy of these species. Variants with coverage below 30 or with a minimum fraction of reads covering a variant (--min_AF) below 0.25 were filtered out. Moreover, we filtered variants to retain only those that passed the filters of all three calling tools, ensuring a collection of high-confidence variants per sample. All variants identified in WT and in the corresponding YPD control treatment, or in the parental for switch treatment strains (ANI for AinF and FLZ for FinA regimens, respectively) were removed from the high-confidence sets (with BCFtools v. 1.15.1^113^, function isec) to distinguish mutations that appeared during the evolution experiment from the pre-existing background genetic variation.

We annotated newly acquired variants for each strain with perSVade, (annotate_small_vars module with Ensembl Variant Effect Predictor v. 100.2^116^. We used NCBI Table 12 and Table 4 (https://www.ncbi.nlm.nih.gov/Taxonomy/Utils/wprintgc.cgi) for Genetic and Mitochondrial Genetic code translation, respectively. In most of the analyses, the mitochondrial chromosome from the *C. parapsilosis* genome (HE605210.1) was excluded unless indicated otherwise.

Protein-altering variants were linked to chromosomal feature information from the *C. parapsilosis* CDC317 vs01-m06-r05 reference genome available on CGD aligning affected genes with either their commonly known gene aliases in *C. parapsilosis* reference genome or, if no alias existed, with the corresponding *Saccharomyces cerevisiae* or *Candida albicans* ortholog name(s) and functional description. For *C. orthopsilosis* and *C. metapsilosis*, we performed gene annotation with eggNOG-mapper v2.1.12^117^ using the eggNOG v5. 0.2 database as a reference^118^.

### Detection of copy number variants and aneuploidies

To infer copy number variants (CNVs) and aneuploidies we used variations in read-depth coverage, as implemented in the perSVade “call_CNVs” module. First, this module calculates relative coverage across genomic windows using bedtools (v2.29.0)^119^ and mosdepth (v0.2.6)^120^. This module corrects for mappability, GC content, as well as distance to telomere for samples affected by the “smiley effect” (where relative coverage correlates negatively with distance to telomere)^109^. After correction PerSVade detects copy-number changes and breakpoints using HMMcopy (v1.32.0)^121^ and AneuFinder (v1.18.0)^122^. We set the –-window_size_CNVcalling parameter to 500 to analyze windows >500 bp, calculating relative read depth compared to the median read depth per gene across all nuclear chromosomes without large duplications. The ‘call_CNVs’ module then generates consensus CNV calls from the two algorithms by selecting the most conservative copy number for each window (in diploid organisms this parameter can be 0 (homozygous loss), 0.5 (heterozygous loss), 1.5 (trisomy), or 2.0 (tetrasomy)). Because the CNV prediction algorithms lack single base pair resolution, the same CNV region may be predicted with slightly different coordinates in different samples. To address this, we concatenated and merged the predicted CNVs in WT and YPD samples for each WT strain using bedtools merge (v2.30.0). This generated a list of background CNVs that were filtered out (with bedtools subtract v2.30.0) from the CNV regions identified in the corresponding evolved strains. This step ensured that only CNVs appearing during drug exposure were retained. After this, we performed additional filtering based on both the predicted relative copy number and the relative coverage (measured as the ratio between the median coverage of the CNV region and the median coverage across the whole genome). We required copy number≥1.5 and relative coverage≥1.3 for duplications and copy number≤0.5 and relative coverage≤0.6 for deletions.

For duplications, CNV regions were filtered out if they were shorter than the gene they affected or if they overlapped less than 80% of the length of the gene. For deletions, regions shorter than 1kb or covering less than 10% of the affected gene were also discarded. We functionally annotated CNVs as described for small variants.

### Analysis of structural rearrangements

To detect and compare structural variants (SVs) we developed and used several modules implemented in perSVade (commit ‘d25e82b’ from 10/07/2025 at https://github.com/Gabaldonlab/perSVade). Parameter selection for SV breakpoint calling was performed with the *optimize_parameters* module for each sample (with default parameters and –-simulation_ploidies set to diploid_hetero with –-keep_simulation_files for further benchmarking). Repeats for each species were generated using the respective genome reference with *infer_repeats* module and used in all downstream steps via –-repeats_file. We evaluated the resulting set of parameters with the *analyze_SV_parameters* module across the simulations and selected the most conservative, high-performing configuration (file more_conservative_parameters_{sample}.json). These parameters were further modified manually in order to reduce the false positives call rate, while ensuring they still outperformed the default parameters (higher F-values), according to the *analyze_SV_parameters* results. The curated JSON files were deposited in the GitHub repository (https://github.com/Gabaldonlab/Cpara_complex_evol). SVs were then called with the call_SVs module using the curated parameters. For SV calling and integration, the repeat file was manually modified to retain only simple and low-complexity entries, which were used as –-repeats_file. For downstream interpretation, we integrated SVs with CNVs calls using the *integrate_SV_CNV_calls* module and annotated them with the *annotate_SVs* module. To identify SVs gained or lost in evolved strains relative to their background samples (WT, YPD samples, and also ANI and FLZ for respective AinF and FinA samples) we used the *integrate_several_samples* module. Variants of the same type were considered the same across samples if they overlapped by >75% and had breakends within 50 bp (--pct_overlap 75, –-tol_bp 50) of each other. To filter the set of changing SVs we prioritised entries with type_background = low_confidence, which leads to more conservative (higher confidence) results. In this setting, “gained” variants lacked support from any raw (unfiltered) gridss breakpoints in ≥1 background, while “lost” variants were present in all backgrounds but had no raw-breakpoint support in the target sample. Finally, to focus on reliable new SVs we analysed only “gained” SVs classified as “real” by CLOVE. All of them belonged to the “tandem duplication” or “deletion” category, so we manually curated them using a custom script that considers the coverage around each event to exclude possible artefacts. A detailed description of such custom analyses can be found in https://github.com/Gabaldonlab/perSVade/wiki/9.-Custom-workflows

### Synteny analysis

Genomic synteny between the three species was assessed by mapping the reference genomes against each other with the long-read mapper minimap (v2.28) using the asm5 preset. To lift the aneuploidy callings between species, aneuploid regions in *C. orthopsilosis* and *C. metapsilosis* were manually delimited in BED files and the corresponding sequences extracted with bedtools (v2.30.0). These extracted sequences were then mapped to the *C. parapsilosis* reference genome with minimap (v2.28) to identify homologous regions. Centromere positions were calculated based on^83^ after establishing chromosome correspondence by cross-mapping.

### Inference of loss of heterozygosity blocks

We inferred LOH blocks in hybrid strains of *C. orthopsilosis* and *C. metapsilosis* with JLOH (v1.1.1)^90^. For *C. orthopsilosis*, given the availability of a reference genome sequence of parental A (Co90-125), we can assume blocks assigned as LOH to the alternative allele represent the allele of parental B (SY36). For *C. metapsilosis*, no parental strain has been identified to date and the available reference genome (BP57) is chimeric. Therefore, alleles could not be assigned to a specific parental strain.

As recommended in the tool’s manual, we used the JLOH “stats” module to determine the Q50 density of SNPs. We run JLOH “extract” with this Q50 density as the –-min-snps-kbp for each sample (except for haploid strains, in which case we used the Q50 of the corresponding YPD strain) and –-min-frac-cov set to 0.7, with other parameters kept as default. We removed background LOH blocks and uncalled regions in WT strains or in the parental of switch treatment strains (ANI for AinF and FLZ for FinA regimens, respectively) from the evolved strains. We did not remove background blocks found within a larger, newly acquired LOH block in the evolved strain, as we assumed the recombination event that generated the new larger block also affected the pre-existing LOH block. We used Bedtools intersect (v2.30.0) to extract from the GFF files the genes that were covered by LOH blocks.

### Statistical analysis

Statistical analyses were conducted in R v4.4.2 using tidyverse v2.0.0 (including dplyr v1.1.4), rstatix v0.7.2, and effectsize v1.0.1. LOH extension was compared using a two-sided Wilcoxon rank-sum test (function *wilcox.test*) on the percentage of acquired LOH over the entire genome. Differences in resistance levels (rAUC in fluconazole or anidulafungin) between clinical and environmental strains within each treatment group were tested using Wilcoxon rank-sum test, with p-values adjusted using the FDR method (*p.adjust*). The same test was used to compare resistance levels (rAUC for ani) between anidulafungin-evolved strains (ANI, FinA) with mutations within and outside known hotspots. For relative fitness (AUC_R_), deviations from 1 were assessed within each non-WT treatment group using two-sided one-sample Wilcoxon signed-rank tests (*wilcox.test*, mu = 1), with FDR correction for multiple testing.. The effect of aneuploidies on relative fitness and differences in resistance levels across species (within each treatment group) were assessed using a Kruskal-Wallis test (*kruskal_test, rstatix v0.7.2*), when significant (p < 0.05), pairwise comparisons were performed using Dunn’s post-hoc test with FDR correction (*dunn_test, p.adjust.method = “BH”*) between evolved strains without aneuploidies, with aneuploidies, and those that lost aneuploidies in switch conditions. Finally, a paired two-sided t-test (*t.test*) was used to compare relative fitness between AinF strains with acquired *MRR1* mutations and corresponding ANI strains without them.

## Data availability

Raw reads from this study are deposited in the NCBI Sequence Read Archive under BioProject PRJNA1280132. Reviewers can access them via https://dataview.ncbi.nlm.nih.gov/object/PRJNA1280132?reviewer=80ba0j7d33ej5721n3l31tmq9l

WT Co_MCO456 was sequenced previously^123^ (SRA ERR321926, BioProject PRJEB4430) and is reused in this study.

## Supporting information

Supplementay statistics

## Acknowledgements

The TG group acknowledges support from the Spanish Ministry of Science and Innovation for grants PID2021-126067NB-I00, CPP2021-008552, PCI2022-135066-2, and PDC2022-133266-I00, cofounded by ERDF “A way of making Europe”; from the Catalan Research Agency (AGAUR) SGR01551; from the European Union’s Horizon 2020 research and innovation programme (ERC-2016-724173); from the Gordon and Betty Moore Foundation (Grant GBMF9742); from the “La Caixa” foundation (Grant LCF/PR/HR21/00737), and from the Instituto de Salud Carlos III (IMPACT Grant IMP/00019 and CIBERINFEC CB21/13/00061-ISCIII-SGEFI/ERDF). JCNR received a Predoctoral Fellowship from the Spanish Ministry of Science and Innovation (grant number PRE2019-088193). ARR received a Predoctoral Fellowship from the Spanish Ministry of Universities with code FPU22/01226. VB has received funding from the European Union’s Horizon 2020 research and innovation programme under the Marie Skłodowska-Curie grant agreement No 945352. Some of the figures were created with BioRender.com.

The authors thank the Functional Genomics Core Facility of IRB Barcelona for their support in the generation of genomic libraries and sequencing. We thank the Scientific and Technological Centers (CCiTUB), Universitat de Barcelona, and Jaume Comas-Rius for their support and advice on flow cytometry technique. We also appreciate all members of the Gabaldón group for their support throughout this project. In particular, Ester Saus, Alicia Benavente and Alicia Barcala for technical support in the management of DNA extractions.

## Author contributions statements

JCNR conceived the project and the experimental design, performed experiments, selection of strains to sequencing, extraction of the DNA, phenotype experiments and analysis, and wrote the original draft. VB and ARR performed the bioinformatic analyses of sequenced strains, including read processing, SNP, CNV and LOH calling, performed statistical analyses of the results and contributed to writing the original draft, with VB focusing on *C. parapsilosis* and ARR in the other species. MÀST developed bioinformatic code to analyse SVs and collaborated on the interpretation of the results. VdO performed bioinformatic analyses including read processing, SNP and LOH calling. MP and IA contributed to perform phenotype experiments and analysis. TG conceived the project, and both the experimental and computational designs, supervised the analysis, acquired funds and wrote the original draft. All authors have read and accepted the manuscript.

## List of Abbreviations

AUC: Area under the curve
AUC_R_: Area under the curve relative to WT
rAUC: Resistance area under the curve
AinF: Anidulafungin-evolved in switch treatment to fluconazole
ANI: Anidulafungin treatment
ANIFLZ: Anidulafungin and fluconazol combined treatment
CFU: Colony forming unit
CNV: Copy number variant
FLZ: Fluconazole treatment
FinA: Fluconazole-evolved in switch treatment to anidulafungin
GoF: Gain of function
LoF: Loss of function
LOH: Loss of heterozygosity
MIC50: Minimum inhibitory concentration
MIC_R: Minimum inhibitory concentration relative to WT
RR: Resistance ratio
SMG: Supra MIC growth
SV: Structural variant
YPD: Yeast extract peptone dextrose treatment
WT: Wild type

