## Supplementary Figures for "Reversible haploidisation and convergent genomic routes to antifungal resistance in the *Candida parapsilosis* species complex"

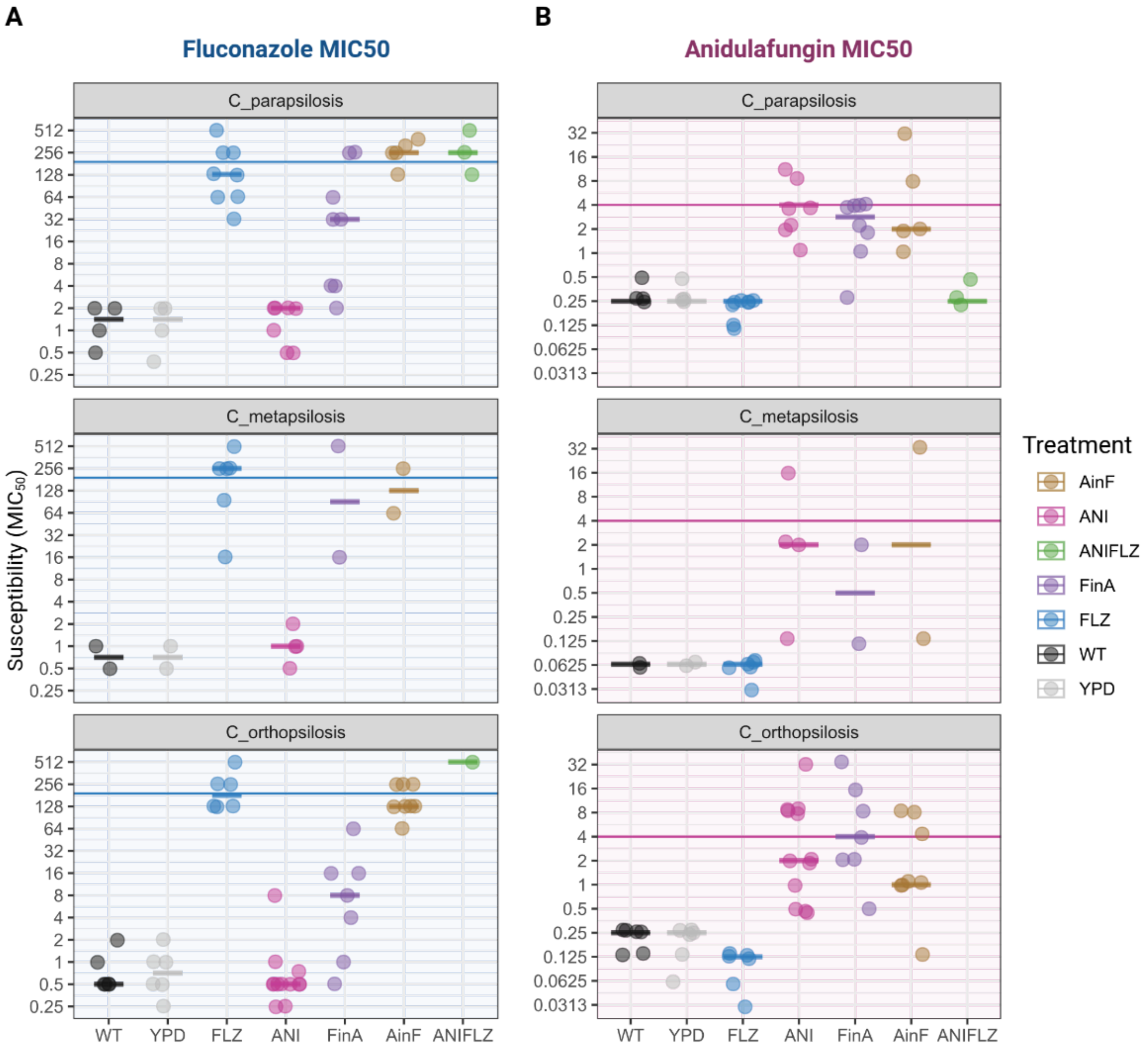


**Supplementary Figure 1. Susceptibility of experimentally evolved strains**. Dot plots showing the Minimum Inhibitory Concentration at 50% ($MIC_{50}$) for each evolved strain of *C. parapsilosis*, *C. metapsilosis* and *C. orthopsilosis* in (**A**) fluconazole and (**B**) anidulafungin. Points represent the median of 4 biological replicates. Bars with the colors of the dots represent the mean between strains for each treatment. The horizontal line across the Y-axis marks the last concentration used in the evolution with (**A**) fluconazole or (**B**) anidulafungin.

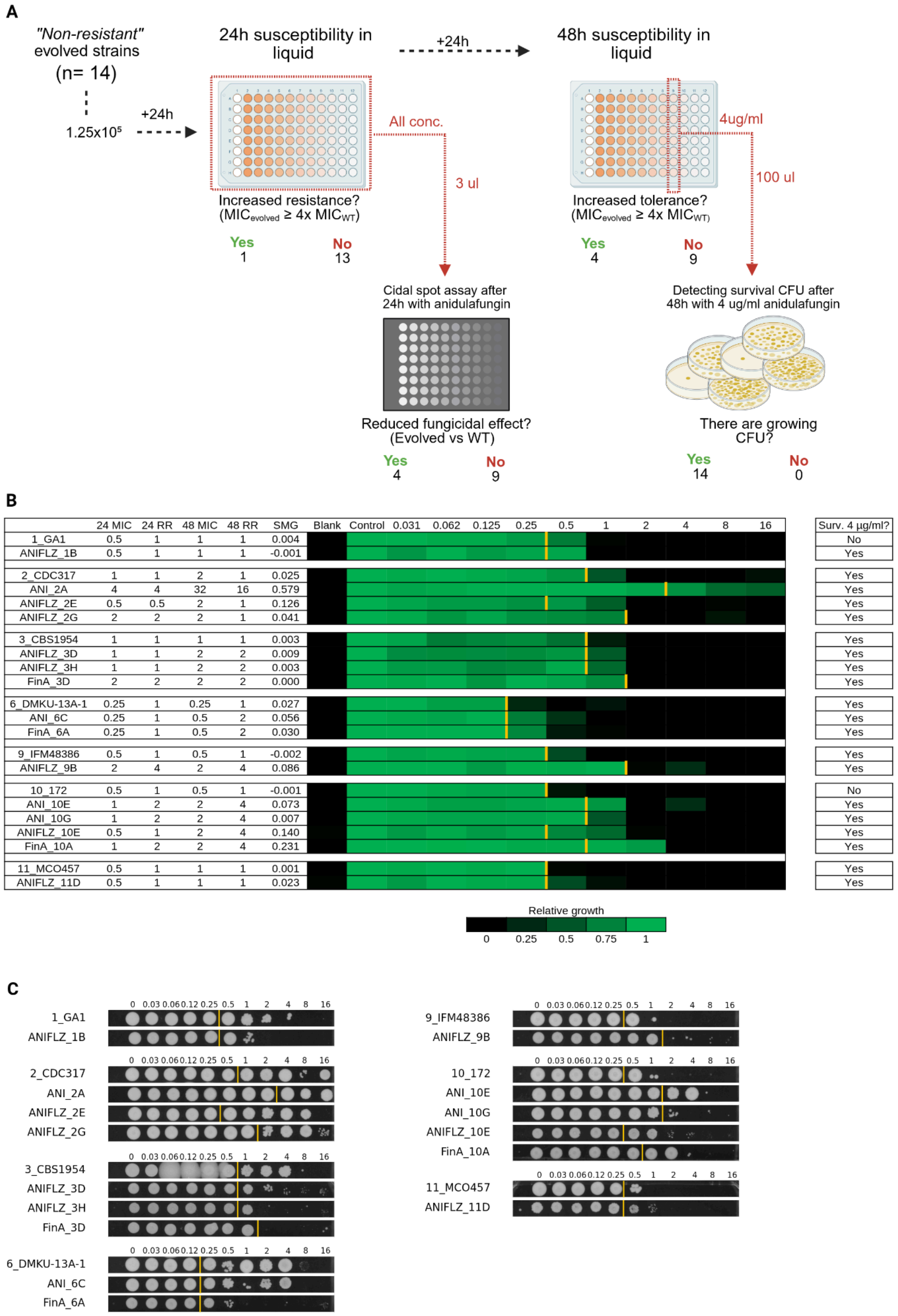
A
"Non-resistant" evolved strains
(n= 14)
1.25x10⁵
+24h
24h susceptibility in liquid
Increased resistance? (MICevolved ≥ 4x MICWT)
Yes 1
No 13
All conc.
3 ul
+24h
48h susceptibility in liquid
4ug/ml
100 ul
Increased tolerance? (MICevolved ≥ 4x MICWT)
Yes 4
No 9
Cidal spot assay after 24h with anidulafungin
Reduced fungicidal effect? (Evolved vs WT)
Yes 4
No 9
Detecting survival CFU after 48h with 4 ug/ml anidulafungin
There are growing CFU?
Yes 14
No 0
B
24 MIC | 24 RR | 48 MIC | 48 RR | SMG | Blank | Control | 0.031 | 0.062 | 0.125 | 0.25 | 0.5 | 1 | 2 | 4 | 8 | 16 | Surv. 4 µg/ml?
1_GA1 | 0.5 | 1 | 1 | 1 | 0.004 | No
ANIFLZ_1B | 0.5 | 1 | 1 | 1 | -0.001 | Yes
2_CDC317 | 1 | 1 | 2 | 1 | 0.025 | Yes
ANI_2A | 4 | 4 | 32 | 16 | 0.579 | Yes
ANIFLZ_2E | 0.5 | 0.5 | 2 | 1 | 0.126 | Yes
ANIFLZ_2G | 2 | 2 | 2 | 1 | 0.041 | Yes
3_CBS1954 | 1 | 1 | 1 | 1 | 0.003 | Yes
ANIFLZ_3D | 1 | 1 | 2 | 2 | 0.009 | Yes
ANIFLZ_3H | 1 | 1 | 2 | 2 | 0.003 | Yes
FinA_3D | 2 | 2 | 2 | 2 | 0.000 | Yes
6_DMKU-13A-1 | 0.25 | 1 | 0.25 | 1 | 0.027 | Yes
ANI_6C | 0.25 | 1 | 0.5 | 2 | 0.056 | Yes
FinA_6A | 0.25 | 1 | 0.5 | 2 | 0.030 | Yes
9_IFM48386 | 0.5 | 1 | 0.5 | 1 | -0.002 | Yes
ANIFLZ_9B | 2 | 4 | 2 | 4 | 0.086 | Yes
10_172 | 0.5 | 1 | 0.5 | 1 | -0.001 | No
ANI_10E | 1 | 2 | 2 | 4 | 0.073 | Yes
ANI_10G | 1 | 2 | 2 | 4 | 0.007 | Yes
ANIFLZ_10E | 0.5 | 1 | 2 | 4 | 0.140 | Yes
FinA_10A | 1 | 2 | 2 | 4 | 0.231 | Yes
11_MCO457 | 0.5 | 1 | 1 | 1 | 0.001 | Yes
ANIFLZ_11D | 0.5 | 1 | 1 | 1 | 0.023 | Yes
Relative growth
0 0.25 0.5 0.75 1
C
0 0.03 0.06 0.12 0.25 0.5 1 2 4 8 16
1_GA1
ANIFLZ_1B
2_CDC317
ANI_2A
ANIFLZ_2E
ANIFLZ_2G
3_CBS1954
ANIFLZ_3D
ANIFLZ_3H
FinA_3D
6_DMKU-13A-1
ANI_6C
FinA_6A
9_IFM48386
ANIFLZ_9B
10_172
ANI_10E
ANI_10G
ANIFLZ_10E
FinA_10A
11_MCO457
ANIFLZ_11D

**Supplementary Figure 2. Characterization of tolerance phenotypes in non-resistant evolved lineages.** (**A**) Schematic representation of the experimental design to investigate survival mechanisms in strains evolved under anidulafungin pressure that did not acquire resistance. Fourteen evolved strains, their respective parental wild-types (WT), and a resistant control (ANI_2A) were subjected to susceptibility testing via broth microdilution with twofold serial dilutions of anidulafungin. After 24 hours, the Minimum Inhibitory Concentration (MIC) and fungicidal activity were assessed. Liquid plates were incubated for an additional 24 hours (48 hours total) to assess delayed susceptibility. Subsequently, viable CFUs were quantified from samples of the maximum evolution concentration (4 µg/ml)(**B**), Summary of liquid assay metrics. The table details MIC values at 24 and 48 hours, the Resistance Ratio, and Supra-MIC Growth (SMG). Graphs display growth relative to the drug-free control (concentration = 0) at 48 hours; yellow markers indicate the 24-hours MIC reference point to visualize supra-MIC growth. The rightmost column denotes the recovery or not of viable CFUs from the 4 µg/ml condition. (C), Spot assays derived from 24-hours susceptibility plate wells (3 µl inocula). The yellow line demarcates the MIC determined in liquid medium at 24 hours. Note: Minor contamination visible in specific spots of strain CBS1954 did not affect data analysis.

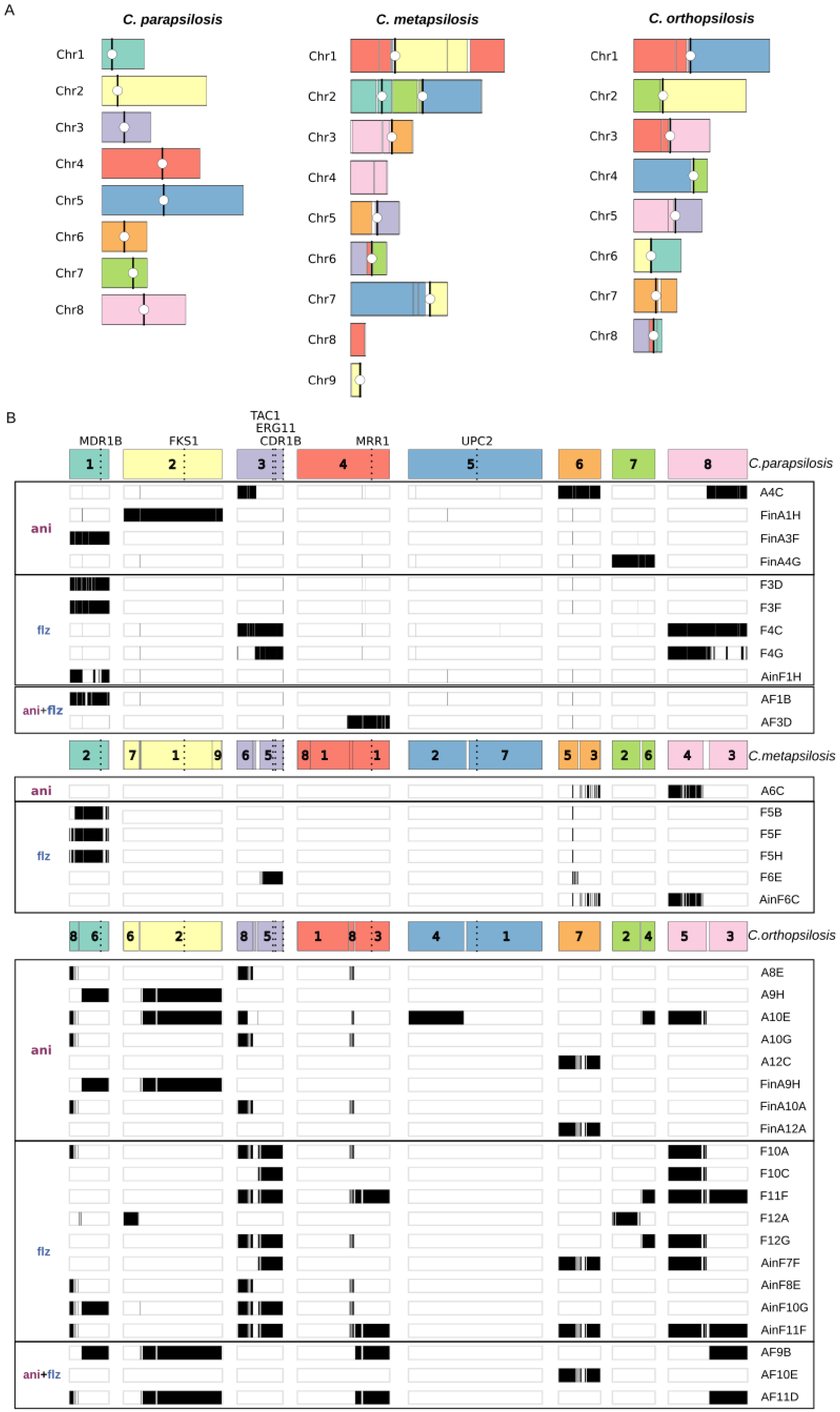
A
C. parapsilosis
C. metapsilosis
C. orthopsilosis
Chr1
Chr2
Chr3
Chr4
Chr5
Chr6
Chr7
Chr8
Chr9
B
MDR1B
FKS1
TAC1
ERG11
CDR1B
MRR1
UPC2
C.parapsilosis
C.metapsilosis
C.orthopsilosis
ani
flz
ani+flz
A4C
FinA1H
FinA3F
FinA4G
F3D
F3F
F4C
F4G
AinF1H
AF1B
AF3D
A6C
F5B
F5F
F5H
F6E
AinF6C
A8E
A9H
A10E
A10G
A12C
FinA9H
FinA10A
FinA12A
F10A
F10C
F11F
F12A
F12G
AinF7F
AinF8E
AinF10G
AinF11F
AF9B
AF10E
AF11D

**Supplementary Figure 3. Drug resistance drives recurrent aneuploidies that affect homologous regions across C. parapsilosis species.** (**A**) Synteny among *C. parapsilosis*, *C. orthopsilosis* and *C. metapsilosis* reference genomes. Syntenic regions share the same colour. Centromeres are indicated as a white dot on a black line. (**B**) Detected aneuploidies in analysed strains, aligning syntenic regions between the three species. In this visualization, reference genomes (in colours) were fragmented and arranged according to their synteny to *C. parapsilosis*. Numbers represent the original chromosome of the synteny block. Dotted lines indicate the location of interest genes, named on top. Black blocks represent aneuploid regions as detected by CNV calling. For *C. parapsilosis*, blocks represent CNV output. For *C. orthopsilosis* and *C. metapsilosis*, blocks represent alignment blocks of manually defined aneuploidies based on CNV output after aligning to the *C. parapsilosis* reference. Strains are grouped according to the drug treatment (anidulafungin, ani; fluconazole, flz; or both drugs, ani+flz).

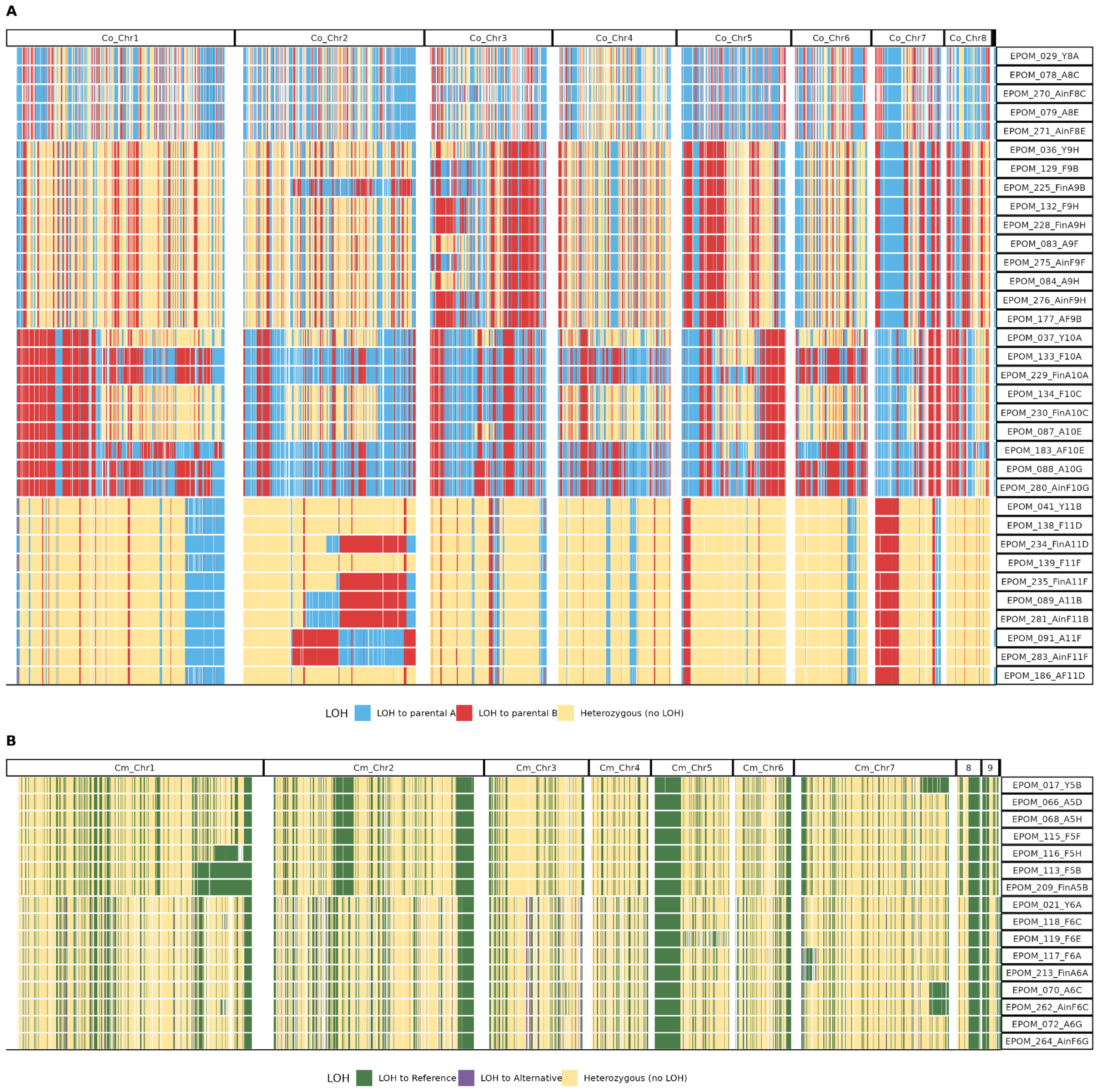


**Supplementary Figure 4. LOH patterns in all evolved hybrid strains of (A) *C. orthopsilosis* and (B) *C. metapsilosis*.** The allele in *C. orthopsilosis* was assigned to parental strains A or B. For *C. metapsilosis*, the allele relative to the chimeric reference genome is reported.

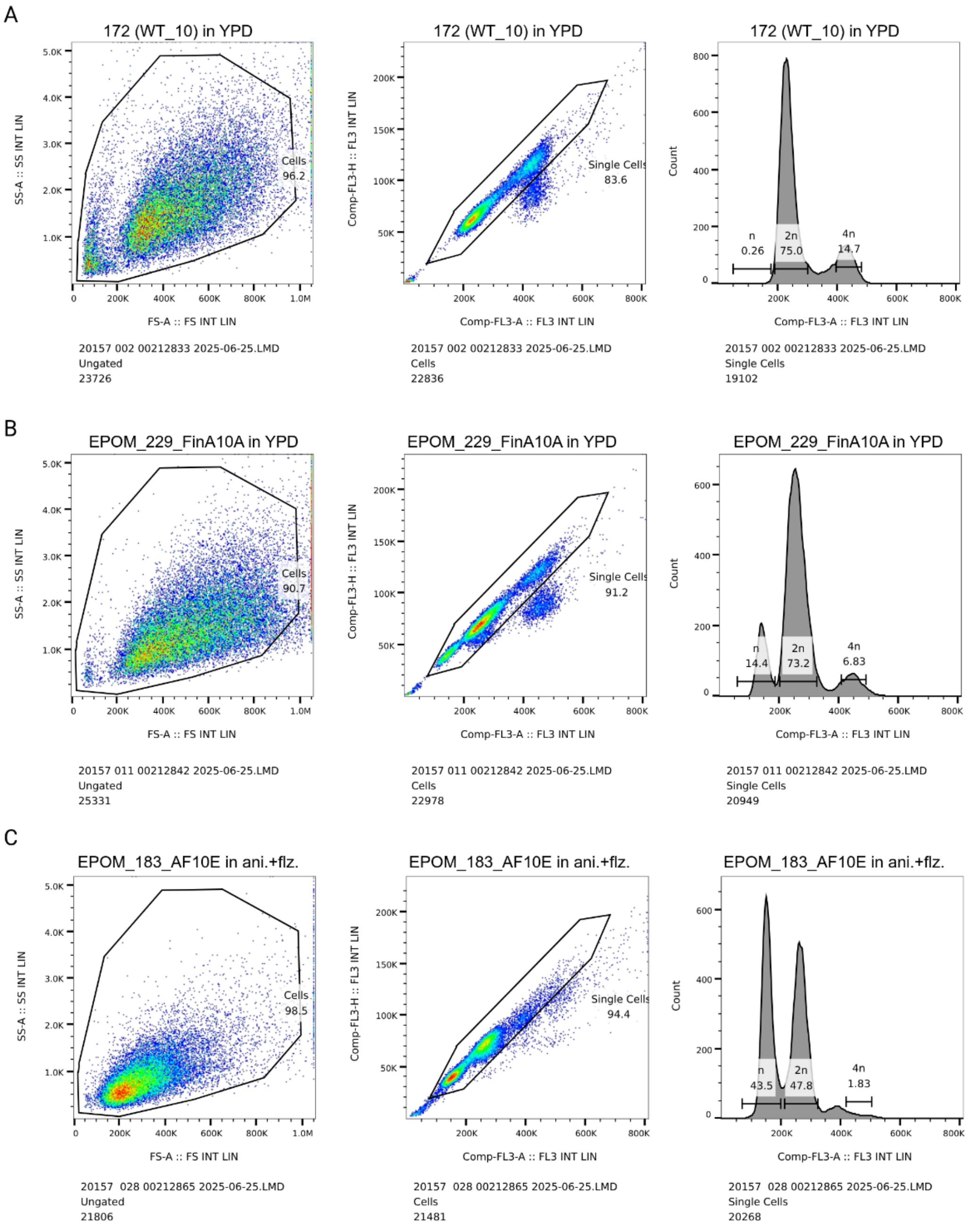


**Supplementary Figure 5. Gating strategy for ploidy determination by flow cytometry.** Representative plots are shown for three distinct samples: (A) a diploid wild-type strain (172_WT_10), (B) an evolved strain with a mixed haploid-diploid population (EPOM_229_FinA10A grown in YPD without drugs), and (C) a predominantly haploid evolved strain (EPOM_183_AF10E). For each sample, the workflow shows gating on the cell population (left), selection of single cells (middle), and the resulting DNA content histogram with ploidy peaks indicated (right).

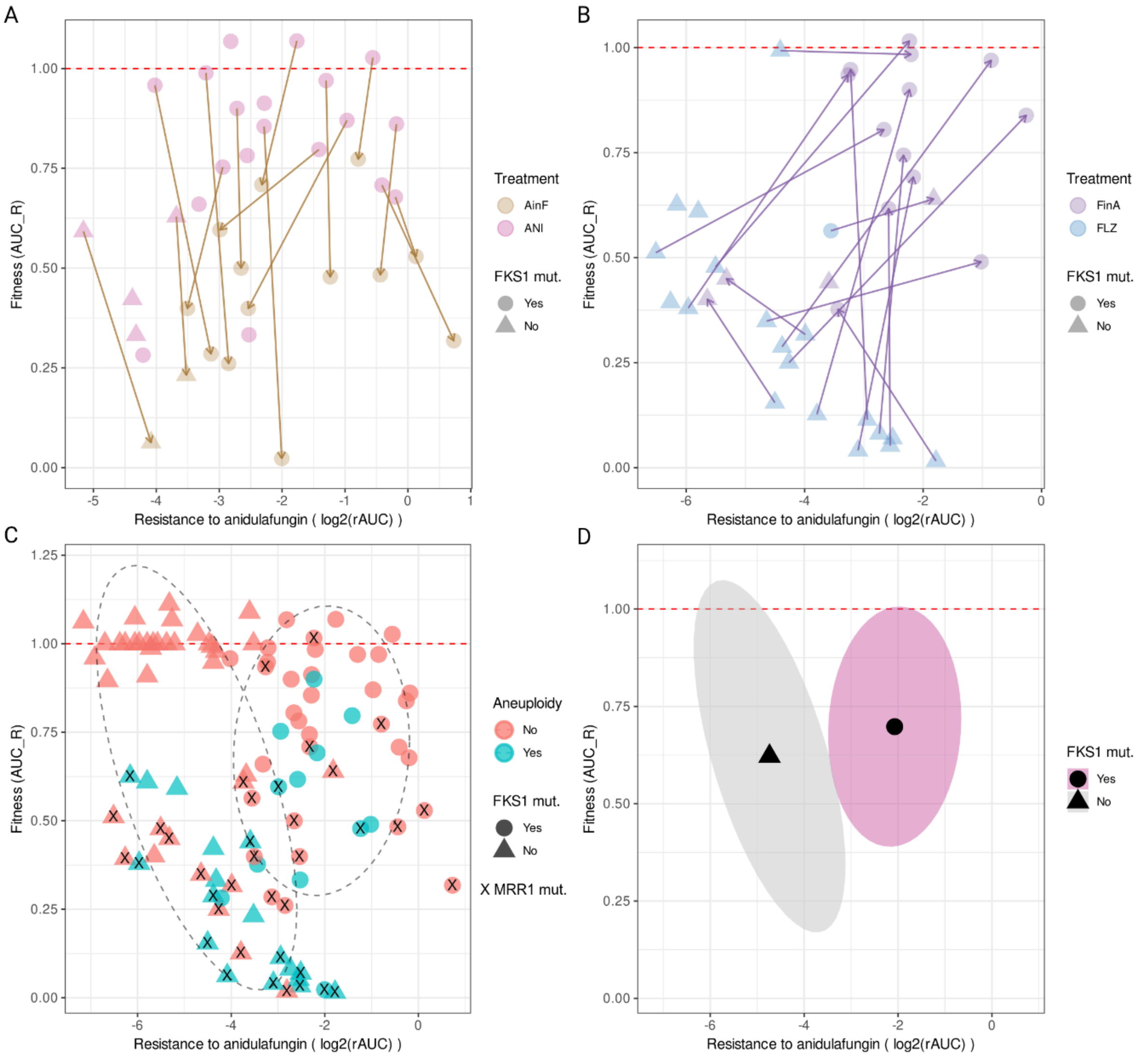


**Supplementary Figure 6. Fitness cost of anidulafungin resistance**. Fitness cost was measured as the ratio between the AUC in YPD of the evolved strain and the AUC in YPD of their WT strain, named as $AUC_R$. The red dotted line indicates the fitness of the WT as a reference, so if it is above, the fitness is increasing, and if it is below, it means that they have decreased their fitness. Resistance to anidulafungin was measured as log2 of the resistance AUC (rAUC) and point shape indicates the presence of missense mutations (Yes) or the absence of mutation (No) in *FKS1*. (A) Fitness ($AUC_R$) of strains evolved in ANI and AinF conditions versus resistance to anidulafungin (log2(rAUC)). Arrows connect paired ANI and its switch evolved AinF strains. (B) Fitness ($AUC_R$) of strains evolved in FLZ and FinA conditions versus resistance to anidulafungin (log2(rAUC)). Arrows connect paired FLZ and its switch evolved FinA strains. (C) Fitness cost associated with anidulafungin resistance and its origins. Dotted ellipses group 50% of samples according to the presence and type of *FKS1* mutations, “X” indicates the strains that also have MRR1 mutations. (D) Dynamics of fitness cost in the development of anidulafungin resistance and later compensatory mutations in switch conditions. Coloured ellipses group 50% samples according to the presence and type of *FKS1* mutations.
