## Supplementay statistics for "Reversible haploidisation and convergent genomic routes to antifungal resistance in the *Candida parapsilosis* species complex"

### Supplementary Note 1: Statistics

#### Drug exposure leads to acquisition of drug resistance and tolerance independently of species or isolation niche

**Resistance levels of species across antifungal treatments.**
For each treatment condition (**treat**) and resistance variable (**var**), the table reports resistance levels by species (**species**) with the sample size (**n**), mean and standard deviation (**mean, sd**), and the median and interquartile range (**median, iqr**).

| treat | var | species | n | mean | sd | median | iqr |
|---|---|---|---|---|---|---|---|
| ANI | ANI_rAUC | C_metapsilosis | 4 | 0.273 | 0.325 | 0.156 | 0.202 |
| ANI | ANI_rAUC | C_orthopsilosis | 12 | 0.257 | 0.253 | 0.141 | 0.311 |
| ANI | ANI_rAUC | C_parapsilosis | 8 | 0.381 | 0.326 | 0.205 | 0.539 |
| ANI | FLZ_rAUC | C_metapsilosis | 4 | 0.006 | 0.002 | 0.006 | 0.002 |
| ANI | FLZ_rAUC | C_orthopsilosis | 12 | 0.040 | 0.036 | 0.026 | 0.067 |
| ANI | FLZ_rAUC | C_parapsilosis | 8 | 0.023 | 0.013 | 0.018 | 0.012 |
| ANIFLZ | FLZ_rAUC | C_orthopsilosis | 1 | 6.721 | | 6.721 | 0.000 |
| ANIFLZ | FLZ_rAUC | C_parapsilosis | 3 | 2.759 | 2.818 | 1.341 | 2.537 |
| AinF | ANI_rAUC | C_metapsilosis | 2 | 0.858 | 1.130 | 0.858 | 0.799 |
| AinF | ANI_rAUC | C_orthopsilosis | 8 | 0.249 | 0.225 | 0.166 | 0.140 |

| | | | | | | | |
|---|---|---|---|---|---|---|---|
| AinF | ANI_rAUC | C_parapsilosis | 5 | 0.435 | 0.412 | 0.249 | 0.439 |
| AinF | FLZ_rAUC | C_metapsilosis | 2 | 0.575 | 0.486 | 0.575 | 0.344 |
| AinF | FLZ_rAUC | C_orthopsilosis | 8 | 0.751 | 0.348 | 0.714 | 0.489 |
| AinF | FLZ_rAUC | C_parapsilosis | 5 | 1.129 | 0.350 | 1.045 | 0.151 |
| FLZ | ANI_rAUC | C_metapsilosis | 6 | 0.069 | 0.069 | 0.041 | 0.098 |
| FLZ | ANI_rAUC | C_orthopsilosis | 6 | 0.049 | 0.036 | 0.044 | 0.025 |
| FLZ | ANI_rAUC | C_parapsilosis | 8 | 0.109 | 0.091 | 0.079 | 0.109 |
| FLZ | FLZ_rAUC | C_metapsilosis | 6 | 0.633 | 0.443 | 0.720 | 0.680 |
| FLZ | FLZ_rAUC | C_orthopsilosis | 6 | 1.609 | 1.369 | 1.381 | 0.799 |
| FLZ | FLZ_rAUC | C_parapsilosis | 8 | 1.009 | 0.843 | 0.834 | 0.866 |
| FinA | ANI_rAUC | C_metapsilosis | 2 | 0.066 | 0.058 | 0.066 | 0.041 |
| FinA | ANI_rAUC | C_orthopsilosis | 7 | 0.358 | 0.278 | 0.223 | 0.366 |
| FinA | ANI_rAUC | C_parapsilosis | 8 | 0.169 | 0.081 | 0.183 | 0.072 |
| FinA | FLZ_rAUC | C_metapsilosis | 2 | 0.672 | 0.854 | 0.672 | 0.604 |
| FinA | FLZ_rAUC | C_orthopsilosis | 7 | 0.100 | 0.153 | 0.049 | 0.033 |
| FinA | FLZ_rAUC | C_parapsilosis | 8 | 0.348 | 0.373 | 0.139 | 0.391 |
| WT | ANI_rAUC | C_metapsilosis | 2 | 0.013 | 0.001 | 0.013 | 0.001 |
| WT | ANI_rAUC | C_orthopsilosis | 6 | 0.020 | 0.005 | 0.020 | 0.007 |
| WT | ANI_rAUC | C_parapsilosis | 4 | 0.040 | 0.034 | 0.032 | 0.038 |

| | | | | | | | |
|---|---|---|---|---|---|---|---|
| WT | FLZ_rAUC | C_metapsilosis | 2 | 0.018 | 0.018 | 0.018 | 0.013 |
| WT | FLZ_rAUC | C_orthopsilosis | 6 | 0.011 | 0.010 | 0.007 | 0.011 |
| WT | FLZ_rAUC | C_parapsilosis | 4 | 0.021 | 0.012 | 0.015 | 0.006 |
| YPD | ANI_rAUC | C_metapsilosis | 2 | 0.009 | 0.002 | 0.009 | 0.002 |
| YPD | ANI_rAUC | C_orthopsilosis | 6 | 0.025 | 0.012 | 0.022 | 0.008 |
| YPD | ANI_rAUC | C_parapsilosis | 4 | 0.045 | 0.030 | 0.044 | 0.027 |
| YPD | FLZ_rAUC | C_metapsilosis | 2 | 0.022 | 0.023 | 0.022 | 0.017 |
| YPD | FLZ_rAUC | C_orthopsilosis | 6 | 0.010 | 0.010 | 0.005 | 0.009 |
| YPD | FLZ_rAUC | C_parapsilosis | 4 | 0.027 | 0.023 | 0.018 | 0.013 |

**Species differences in resistance levels within each treatment condition.**
For each treatment condition (**treat**) and resistance variable (**var**), the table reports comparison across species using the Kruskal-Wallis test, including the test statistic (**stat,** $\chi^2$), degrees of freedom (**df**), and p-value (**p_value**). Effect size is reported as $\eta^2[H]$ (**es**).

| **treatment** | **var** | **stat** | **df** | **p_value** | **es** | **p_adj** |
|---|---|---|---|---|---|---|
| FinA | FLZ_rAUC | 4.672 | 2 | 9.67E-02 | 0.191 | **3.42E-01** |
| YPD | ANI_rAUC | 4.359 | 2 | 1.13E-01 | 0.262 | **3.42E-01** |
| ANI | FLZ_rAUC | 4.065 | 2 | 1.31E-01 | 0.098 | **3.42E-01** |
| AinF | FLZ_rAUC | 3.656 | 2 | 1.61E-01 | 0.138 | **3.42E-01** |
| FinA | ANI_rAUC | 3.571 | 2 | 1.68E-01 | 0.112 | **3.42E-01** |
| ANIFLZ | FLZ_rAUC | 1.800 | 1 | 1.80E-01 | 0.400 | **3.42E-01** |

| | | | | | | |
|---|---|---|---|---|---|---|
| WT | ANI_rAUC | 3.058 | 2 | 2.17E-01 | 0.118 | **3.42E-01** |
| FLZ | FLZ_rAUC | 2.970 | 2 | 2.26E-01 | 0.057 | **3.42E-01** |
| YPD | FLZ_rAUC | 2.878 | 2 | 2.37E-01 | 0.098 | **3.42E-01** |
| WT | FLZ_rAUC | 1.808 | 2 | 4.05E-01 | -0.021 | **5.27E-01** |
| FLZ | ANI_rAUC | 1.562 | 2 | 4.58E-01 | -0.026 | **5.41E-01** |
| ANI | ANI_rAUC | 1.354 | 2 | 5.08E-01 | -0.031 | **5.50E-01** |
| AinF | ANI_rAUC | 0.585 | 2 | 7.46E-01 | -0.118 | **7.46E-01** |

**Resistance levels of clinical versus environmental isolates across antifungal treatments.**
For each treatment (**treat**) and resistance variable (**var**) (FLZ_rAUC, ANI_rAUC), the table reports the sample size for for clinical (**g1**) and environmental (**g2**) isolates (**n1**, **n2**), with corresponding group's median and interquartile range (**med1/iqr1**, **med2/iqr2**). Group differences are reported as the estimated effect (**est**) with 95% confidence interval (**CI_l-CI_h**), alongside the Wilcoxon rank-sum test statistics (**stat,** W) and p-value (**p_value**). Effect size reported as rank-biserial r (**es**) with its 95% confidence interval (**es_CI_l-es_CI_h**). Raw p-values were adjusted for multiple comparisons using FDR (**p_adj**).

| treat | var | g1 | g2 | n1 | n2 | med1 | iqr1 | med2 | iqr2 | est | CI_l | CI_h | stat | p_value | es | es_CI_l | es_CI_h | p_adj |
|---|---|---|---|---|---|---|---|---|---|---|---|---|---|---|---|---|---|---|
| **ANI** | FLZ_rAUC | clin | env | 16 | 8 | 0.021 | 0.055 | 0.007 | 0.007 | 0.014 | 0.003 | 0.059 | 104.5 | 1.43E-02 | 0.633 | 0.241 | 0.847 | **1.85E-01** |
| **FinA** | ANI_rAUC | clin | env | 11 | 6 | 0.213 | 0.255 | 0.130 | 0.149 | 0.099 | -0.010 | 0.388 | 50 | 9.73E-02 | 0.515 | -0.021 | 0.821 | **6.32E-01** |
| **WT** | FLZ_rAUC | clin | env | 8 | 4 | 0.017 | 0.021 | 0.010 | 0.010 | 0.010 | -0.008 | 0.027 | 24.5 | 1.73E-01 | 0.531 | -0.129 | 0.865 | **6.59E-01** |
| **YPD** | FLZ_rAUC | clin | env | 8 | 4 | 0.017 | 0.025 | 0.009 | 0.009 | 0.008 | -0.008 | 0.044 | 24 | 2.03E-01 | 0.500 | -0.170 | 0.854 | **6.59E-01** |

| | | | | | | | | | | | | | | | | | | |
|---|---|---|---|---|---|---|---|---|---|---|---|---|---|---|---|---|---|---|
| **FLZ** | ANI_rAUC | clin | env | 12 | 8 | 0.048 | 0.055 | 0.090 | 0.117 | -0.031 | -0.118 | 0.025 | 35 | 3.35E-01 | -0.271 | -0.668 | 0.246 | **8.71E-01** |
| **YPD** | ANI_rAUC | clin | env | 8 | 4 | 0.026 | 0.023 | 0.014 | 0.016 | 0.009 | -0.025 | 0.040 | 21 | 4.45E-01 | 0.313 | -0.378 | 0.780 | **8.71E-01** |
| **WT** | ANI_rAUC | clin | env | 8 | 4 | 0.020 | 0.009 | 0.016 | 0.023 | 0.003 | -0.067 | 0.017 | 20 | 5.52E-01 | 0.250 | -0.435 | 0.752 | **8.71E-01** |
| **FinA** | FLZ_rAUC | clin | env | 11 | 6 | 0.068 | 0.369 | 0.135 | 0.023 | -0.065 | -0.117 | 0.333 | 27 | 5.80E-01 | -0.182 | -0.650 | 0.386 | **8.71E-01** |
| **ANI** | ANI_rAUC | clin | env | 16 | 8 | 0.187 | 0.308 | 0.137 | 0.282 | 0.049 | -0.125 | 0.275 | 73 | 6.03E-01 | 0.141 | -0.344 | 0.566 | **8.71E-01** |
| **AinF** | FLZ_rAUC | clin | env | 11 | 4 | 0.888 | 0.496 | 0.770 | 0.587 | 0.122 | -0.662 | 0.657 | 24 | 8.45E-01 | 0.091 | -0.531 | 0.649 | **1.00E+00** |
| **FLZ** | FLZ_rAUC | clin | env | 12 | 8 | 0.957 | 0.821 | 0.777 | 1.204 | 0.115 | -1.267 | 0.653 | 51 | 8.47E-01 | 0.063 | -0.436 | 0.531 | **1.00E+00** |
| **AinF** | ANI_rAUC | clin | env | 11 | 4 | 0.172 | 0.205 | 0.426 | 0.867 | -0.027 | -1.457 | 0.311 | 21 | 9.48E-01 | -0.045 | -0.622 | 0.563 | **1.00E+00** |
| **ANIFLZ** | FLZ_rAUC | clin | env | 2 | 2 | 3.827 | 2.895 | 3.673 | 2.332 | -0.271 | -5.073 | 5.380 | 2 | 1.00E+00 | 0.000 | -0.852 | 0.852 | **1.00E+00** |

**Resistance levels of hybrid versus non-hybrid strains of *C. orthopsilosis*.**

Analysing strains based on the last adaptation treatment and taking their resistance to that drug (**var,** FLZ_rAUC, ANI_rAUC), the table reports the sample size for for hybrid (**g1**) and non-hybrid (**g2**) isolates (**n1**, **n2**), with corresponding group's median and interquartile range (**med1/iqr1**, **med2/iqr2**). Group differences are reported as the estimated effect (**est**) with 95% confidence interval (**CI_low-CI_high**), alongside the Wilcoxon rank-sum test statistics (**W**) and p-value (**p_value**). Effect size reported as rank-biserial (**r**).

| var | g1 | g2 | n1 | n2 | med1 | iqr1 | med2 | iqr2 | est | W | CI_low | CI_high | r | p_value |
|---|---|---|---|---|---|---|---|---|---|---|---|---|---|---|
| ANI-rAUC | Hybrid | Parental | 14 | 5 | 0.304 | 0.361 | 0.267 | 0.145 | 0.052 | 42 | -0.175 | 0.352 | 0.149 | 0.5593 |
| FLZ-rAUC | Hybrid | Parental | 11 | 3 | 0.123 | 0.121 | 0.314 | 0.325 | -0.097 | 15 | -3.596 | 0.885 | 0.062 | 0.8846 |

### Anidulafungin resistance is mediated by a broad spectrum of *FKS1* variants

**Anidulafungin resistance levels in strains with *FKS1* mutation within vs outside hotspot regions.**
For the anidulafungin resistance levels (ANI_rAUC), the table reports the sample size (**n**), median and interquartile range (**med, iqr**), mean and standard deviation (**mean, sd**), and the Shapiro-Wilk test p-value (**Shapiro_p**) for each comparison group (Hotspot: No vs Yes).

| Hotspot | n | med | iqr | mean | sd | Shapiro_p |
|---|---|---|---|---|---|---|
| No | 9 | 0.167 | 0.091 | 0.213 | 0.208 | 0.000198 |
| Yes | 23 | 0.217 | 0.347 | 0.345 | 0.263 | 0.00265 |

**Species differences in resistance levels within each treatment condition.**
For anidulafungin resistance levels (ANI_rAUC), the table reports the two-group comparison (Wilcoxon rank-sum test) between strains with *FKS1* mutations outside (**g1**) vs within (**g2**) hotspot regions, including corresponding sample sizes (**n1, n2**), the estimated effect (**est**) with 95% confidence interval (**CI_l-CI_h**), the test statistic (**stat**, W) and p-value (**p_value**), and rank-biserial r effect size (**es**) with its 95% confidence interval (**es_CI_l-es_CI_h**).

| var | g1 | g2 | n1 | n2 | est | CI_l | CI_h | stat | p_value | es | es_CI_l | es_CI_h |
|---|---|---|---|---|---|---|---|---|---|---|---|---|
| ANI_rAUC | No | Yes | 9 | 23 | -0.075 | -0.298 | 0.015 | 66 | **1.21E-01** | -0.362 | -0.681 | 0.072 |

### LOH favors resistance-associated alleles in hybrid strains

**Acquired LOH in hybrid strains between treatment conditions and species**

Acquired LOH was calculated for each hybrid strain as the ratio between the sum of all LOH blocks predicted by JLOH against the total length of the reference genome. These statistics were calculated with bedtools based on the BED files generated by JLOH extract. Haploidized and non-hybrid strains were excluded from this analysis.A Wilcoxon signed-rank test was used to compare groups. The first row is the comparison of YPD-evolved (**g1**) and drug-evolved (**g2**) strains. The second row reports the comparison of *C. metapsilosis* strains (**g1**) and *C. orthopsilosis* strains (**g2**). The table reports each group's size (**n**), median (**med**) and interquartile range (**iqr**). The estimate (**est**) with 95% confidence interval (**CI_low-CI_high**), alongside the Wilcoxon rank-sum test statistics (**W**), the rank-biserial r (**r**) and p-value (**p_value**) are reported.

| var | g1 | g2 | n1 | n2 | med1 | iqr1 | med2 | iqr2 | est | W | CI_low | CI_high | r | p_value |
|---|---|---|---|---|---|---|---|---|---|---|---|---|---|---|
| LOH | YPD | Evol | 6 | 39 | 0.020 | 0.025 | 0.046 | 0.048 | -0.018 | 63 | -0.048 | 9.8e-4 | 0.269 | 0.07357 |
| LOH | Meta_evol | Ortho_evol | 14 | 25 | 0.022 | 0.026 | 0.059 | 0.081 | -0.028 | 110 | -0.053 | 9.6e-4 | 0.305 | 0.05832 |

### Fitness costs are widespread in drug resistance

#### Fitness costs associated with antifungal adaptation

**Fitness costs associated with antifungal adaptation.**
For each treatment (**treat**), number of evolved strains (**n**) and the distribution of relative fitness (**AUC_R**) summarised as median (**med**) and interquartile range (**iqr**). Fitness deviation from the WT strains was tested within each group using a two-sided one-sample Wilcoxon signed-rank test against a null median of 1 (*wilcox.test*, mu = 1), with the corresponding 95% confidence interval for the median difference (**CI_l**, **CI_h**) and test statistic (**stat**, W). Effect sizes are reported as rank-biserial r (**es**) and p-values (**p_value**) were adjusted for multiple testing using FDR (**p_adj**).

| **treat** | **n** | **med** | **iqr** | **CI_l** | **CI_h** | **stat** | **p_value** | **es** | **p_adj** |
|---|---|---|---|---|---|---|---|---|---|
| **FLZ** | 22 | 0.269 | 0.384 | 0.160 | 0.390 | 0 | 4.30E-05 | -1.00 | **2.58E-04** |
| **ANI** | 24 | 0.826 | 0.306 | 0.661 | 0.885 | 15 | 1.22E-04 | -0.90 | **3.65E-04** |
| **FinA** | 17 | 0.744 | 0.445 | 0.597 | 0.857 | 1.5 | 4.20E-04 | -0.98 | **7.22E-04** |
| **AinF** | 16 | 0.399 | 0.254 | 0.252 | 0.503 | 0 | 4.81E-04 | -1.00 | **7.22E-04** |
| **ANIFLZ** | 8 | 0.009 | 0.019 | 0.003 | 0.310 | 0 | 1.43E-02 | -1.00 | **1.71E-02** |
| **YPD** | 12 | 1.007 | 0.112 | 0.960 | 1.067 | 45 | 6.66E-01 | 0.15 | **6.66E-01** |

#### Fitness costs associated with presence of aneuploidies

**Descriptives for each aneuploidy group.**
For each aneuploidy group (No, Rev, Yes), sample size (**n**), median relative fitness (**med**), and interquartile range (**iqr**) are indicated.

| **Aneuploidy_rev** | **n** | **med** | **iqr** |
|---|---|---|---|
| No | 69 | 0.909 | 0.436 |
| Rev | 8 | 0.655 | 0.361 |
| Yes | 34 | 0.194 | 0.438 |

**Kruskal-Wallis test of relative fitness across aneuploidy groups.**
Differences in relative fitness (AUC_R) across aneuploidy groups were tested using a Kruskal-Wallis test. Test statistic (**stat**), degrees of freedom (**df**), and p-value (**p_value**) are indicated. Effect size is reported as $\eta^2[H]$ (**es**).

| **group_var** | **stat** | **df** | **p_value** | **es** |
|---|---|---|---|---|
| Aneuploidy_rev | 42.917 | 2 | 4.79E-10 | 0.379 |

**Dunn's post-hoc pairwise comparison.**
Pairwise differences between aneuploidy groups were tested using Dunn's test, **comp** indicates the compared groups and **stat** is the z statistic. Raw p-values (**p_value**) were adjusted for multiple testing using FDR (**p_adj**).

| **group_var** | **comp** | **stat** | **p_value** | **p_adj** |
|---|---|---|---|---|
| Aneuploidy_rev | No vs Yes | -6.551 | 5.71E-11 | **1.71E-10** |
| Aneuploidy_rev | Rev vs Yes | -2.319 | 2.04E-02 | **3.06E-02** |
| Aneuploidy_rev | No vs Rev | -1.236 | 2.17E-01 | **2.17E-01** |

#### Fitness costs associated with acquired *MRR1* mutations

**Descriptive statistics for ANI and AinF strains with aquired *MRR1* mutations.**
For each treatment (**treat**), the sample size (**n**) and summary statistics of relative fitness (AUC_R), including mean (**mean**), standard deviation (**sd**), median (**med**), and Shapiro-Wilk p-values (Shapiro_p) are indicated.

| treat | n | mean | sd | med | Shapiro_p |
|---|---|---|---|---|---|
| ANI | 15 | 0.830 | 0.176 | 0.861 | 5.08E-01 |
| AinF | 15 | 0.388 | 0.233 | 0.399 | 6.08E-01 |

**Summary of within-pair fitness differences.**
The table reports the distribution of paired relative fitness (AUC_R) differences (AinF - ANI) across **n_pairs**, including **mean_diff**, **sd_diff**, **median_diff**, and **iqr_diff**.

| n_pairs | mean_diff | sd_diff | median_diff | iqr_diff |
|---|---|---|---|---|
| 15 | -0.442 | 0.189 | -0.4 | 0.154 |

**Paired t-test comparing relative fitness (AUC_R) between AinF and ANI strains.**
Relative fitness (AUC_R) was compared within pairs using a two-sided paired t-test. The table reports the mean paired difference (**est**) with 95% confidence interval (**CI_l**, **CI_h**), test statistic (**stat**), degrees of freedom (**df**), p-value (**p_value**), and effect size calculated as Cohen's d (**es**) with its confidence interval (**es_CI_l**, **es_CI_h**).

| est | CI_l | CI_h | stat | df | p_value | es | es_CI_l | es_CI_h |
|---|---|---|---|---|---|---|---|---|
| -0.442 | -0.547 | -0.337 | -9.049 | 14 | 3.17E-07 | -2.337 | -3.320 | -1.331 |
